# Metabolism of glucose activates TORC1 through multiple mechanisms in *Saccharomyces cerevisiae*

**DOI:** 10.1101/2022.03.25.485766

**Authors:** Mohammad Alfatah, Liang Cui, Corinna Jie Hui Goh, Jin Huei Wong, Jacqueline Lewis, Wei Jie Poh, Prakash Arumugam

## Abstract

Target of Rapamycin Complex 1 (TORC1) is a conserved eukaryotic protein complex that links the presence of nutrients with cell growth. In *Saccharomyces cerevisiae*, TORC1 activity is positively regulated by the presence of amino acids and glucose in the medium. However, mechanisms underlying nutrient-induced TORC1 activation remain poorly understood. By utilizing a TORC1 activation assay, we demonstrate that differential metabolism of glucose activates TORC1 through three distinct pathways in yeast. The first ‘canonical Rag GTPase-dependent pathway’ requires conversion of glucose to fructose 1,6-bisphosphate which activates TORC1 via the Rag GTPase heterodimer Gtr1^GTP^/Gtr2^GDP^. The second ‘non-canonical Rag GTPase-dependent pathway’ requires conversion of glucose to glucose 6-phosphate which activates TORC1 via Gtr1^GTP^/Gtr2^GTP^. The third ‘Rag GTPase-independent pathway’ requires complete glycolysis and vacuolar ATPase reassembly for TORC1 activation. Glucose-induced TORC1 activation can be uncoupled from glucose-induced AMPK inactivation. We have established a roadmap to deconstruct the link between glucose metabolism and TORC1 activation.

## Introduction

Target of Rapamycin Complex 1 (TORC1) is a master regulator of eukaryotic cell growth, proliferation, and metabolism. TORC1 signalling pathway controls cellular growth through activation of anabolic processes such as protein, lipid, and nucleotide synthesis, and repression of catabolic processes such as autophagy and proteasomal activity. Dysregulation of TORC1 signalling has been implicated in the diverse set of common human diseases, including cancer, neurological disorders, obesity and diabetes.

TORC1 is a serine/threonine protein kinase belonging to the phosphatidylinositol 3-kinase - (PI3K) related family of lipid kinases. The *Saccharomyces cerevisiae* TORC1 is composed of four subunits Tor1 or Tor2 (the catalytic subunit), Kog1 (ortholog of mammalian raptor), Lst8 (synthetic Lethal with Sec Thirteen) and Tco89 (Gonzalez and Hall, 2017). TORC1 in yeast forms a lozenge-shaped dimer that contains two copies of each of the 4 subunits. Human mTORC1’s composition is similar to its yeast counterpart but lacks Tco89.

TORC1 is regulated by the conserved Rag-family of small guanosine triphosphatases (GTPases) namely the Gtr1 (RagA/B) and Gtr2 (RagC/D) proteins in yeast (Powis and De Virgilio, 2016). Gtr1 and Gtr2 heterodimerise and localize to vacuolar membranes *via* an interaction with the EGO (Ego1–Ego2–Ego3) complex (Binda et al., 2010). Amino acid sufficiency promotes the active conformation of the Gtr1/Gtr2 heterodimer in which Gtr1 and Gtr2 are loaded with GTP and GDP respectively (Panchaud et al., 2013). Active Gtr1^GTP/Gtr2GDP^ heterodimer binds to Kog1 and activates TORC. Mammalian RagA/B and RagC/D GTPases bind to the lysosomal Ragulator complex composed of five subunit proteins (LAMTOR1-5) and sense the availability of amino acids (Saxton and Sabatini, 2017). In the presence of amino acids (particularly leucine and arginine), RagA/B and RagC/D GTPases activate mTORC1 by recruiting it to the lysosomes for activation by the lysosomal Rheb GTPase. Notably, mTORC1 but not the budding yeast TORC1 is regulated by the Rheb GTPase. Moreover, budding yeast TORC1 localizes constitutively to the vacuoles unlike mTORC1 which requires Rag-GTPases for its lysosomal localization.

Nucleotide-binding status of Gtr1 and Gtr2 is tightly regulated by conserved GAPs (GTPase-activating proteins) and GEFs (guanine exchange factors). SEACIT (Iml1-Npr2-Npr3) and Lst4-Lst7 complexes act as GAPs for Gtr1 and Gtr2, respectively (Nicastro et al., 2017). Activity of SEACIT is negatively regulated by SEACAT (Sea2, Sea3 and Sea4) complex in yeast (Panchaud et al., 2013). Mammalian counterparts for SEACIT and SEACAT are GATOR1 and GATOR2 complexes respectively. The budding yeast vacuolar protein Vam6 has been proposed to be the GEF for Gtr1 (Panchaud et al., 2013). Vam6 is a component of the conserved HOPS complex involved in homotypic vacuolar fusion and vacuolar protein sorting. Identity of the GEF for Gtr2 remains unknown.

It is intriguing how nutrients regulate TORC1 activity in eukaryotic cells. Somewhat unexpectedly, this process is better understood in mammalian cells in comparison to yeast. In the absence of amino acids, sestrins bind and inhibit mammalian GATOR2 (Chantranupong et al., 2014; Saxton et al., 2016a; Saxton et al., 2016b; Wolfson et al., 2016). This frees up GATOR1 which activates GTP hydrolysis on Rag A/B and inhibits mTORC1. Binding of amino acids like leucine and arginine to sestrins inhibits its interaction with GATOR2 and activates mTORC1. Apart from one report showing that the leucyl-tRNA synthetase (Cdc60) acts as a GEF for Gtr1 in a leucine-dependent manner (Bonfils et al., 2012), there is very little progress in understanding how nutrients activate TORC1 in *Saccharomyces cerevisiae*.

TORC1 activity in yeast is highly sensitive to presence of glucose (Alfatah et al., 2021; Hughes Hallett et al., 2014, 2015; Urban et al., 2007). Availability of glucose regulates disassembly/reassembly of vacuolar ATPase (V-ATPase) (Kane and Smardon, 2003). V-ATPase was proposed to activate TORC1 in response to glucose and amino acids via Gtr1 (Dechant et al., 2014). Specifically, in response to glucose but not nitrogen starvation, TORC1 was reported to disassemble, with its Raptor subunit Kog1 relocating to vacuolar edge to form Kog1 bodies (Hughes Hallett et al., 2015). Formation of Kog1 bodies was driven by Snf1-AMPK mediated phosphorylation of Kog1 at Ser 491/494 and two adjacent prion motifs (Hughes Hallett et al., 2015). When budding yeast cells were subjected to glucose starvation, TORC1 was reported to form catalytically inactive cylindrical structures called as TOROIDs (TORC1 Oligomerised in Inhibited Domain) perched on the vacuolar membrane (Prouteau et al., 2017). While the inactive Gtr1^GDP^/Gtr2^GTP^ promotes TOROID formation, the active Gtr1^GTP^/Gtr2^GDP^ antagonizes their formation (Prouteau et al., 2017). Unlike observed for Kog1 bodies, the Snf1/AMPK did not regulate TOROID formation (Prouteau et al., 2017).

We recently showed that glucose activates TORC1 via Rag GTPase-dependent and Rag GTPase-independent mechanisms (Alfatah et al., 2021). In this study, we have combined genetics with targeted metabolite analysis to determine the extent of glucose metabolism and the upstream TORC1 regulators required for glucose-induced TORC1 activation. We show that glucose activates TORC1 through three distinct pathways. In the first pathway, formation of fructose 1,6-bisphosphate activates TORC1 via the canonical pathway dependent on Gtr1^GTP^/Gtr2^GDP^. In the second pathway, formation of glucose 6-phosphate activates TORC1 via a hitherto undescribed non-canonical pathway involving Gtr1^GTP^/Gtr2^GTP^. Complete glycolysis of glucose to pyruvate activates TORC1 *via* the third Rag GTPase-independent pathway and this requires vacuolar ATPase reassembly and activity. Furthermore, we show that glucose-induced regulation of TORC1 and AMPK activities can be uncoupled from each other indicating that they occur via independent mechanisms.

## Results

### Hexokinase-mediated phosphorylation of glucose is essential for glucose-induced TORC1 activation

We had previously demonstrated that glucose is sufficient to activate TORC1 via Rag GTPase-dependent and Rag GTPase-independent mechanisms in *Saccharomyces cerevisiae* (Alfatah et al., 2021). In the glucose-induced TORC1 activation assay, log phase yeast cells are subjected to complete nutrient starvation by incubating them in water for 1 hour. Glucose is then added to starved cultures and TORC1 activity is assessed by monitoring phosphorylation of its substrate Sch9 (Alfatah et al., 2021). Glucose could either directly activate TORC1 or it might have to be metabolized to activate TORC1. We first tested whether glucose is metabolized by yeast cells during the conditions of our glucose-induced TORC1 activation assay. We performed glucose-induced TORC1 activation assay with wild type and *gtr1Δ* cells and measured the relative levels of glycolytic intermediates by LC-MS (Figure 1A). In starved cells, lower glycolytic intermediates such as phosphoenolpyruvate (PEP) and 3-Phosphoglycerate (3-PG) accumulated (Figure 1A) which is consistent with a previous study (Xu et al., 2012). Upon addition of 2% glucose to starved cells, the relative molar levels of glucose 6-phosphate, fructose 6-phosphate, fructose 1,6-bisphosphate and G3P/DHAP increased and the relative molar levels of PEP and 2PG/3PG decreased (Figure 1A). Moreover, the kinetics of glycolysis in wild type and *gtr1Δ* cells were indistinguishable (Figure 1A). Notably, TORC1 activation in *gtr1Δ* cells was delayed by about 10 minutes in comparison to wild type cells (Figure 1B). These results demonstrate that glucose undergoes glycolysis in yeast cells during the conditions of the glucose-induced TORC1 activation assay. In addition, delayed TORC1 activation in *gtr1*Δ cells is not due to slower rate of glycolysis.

**Figure 1.**
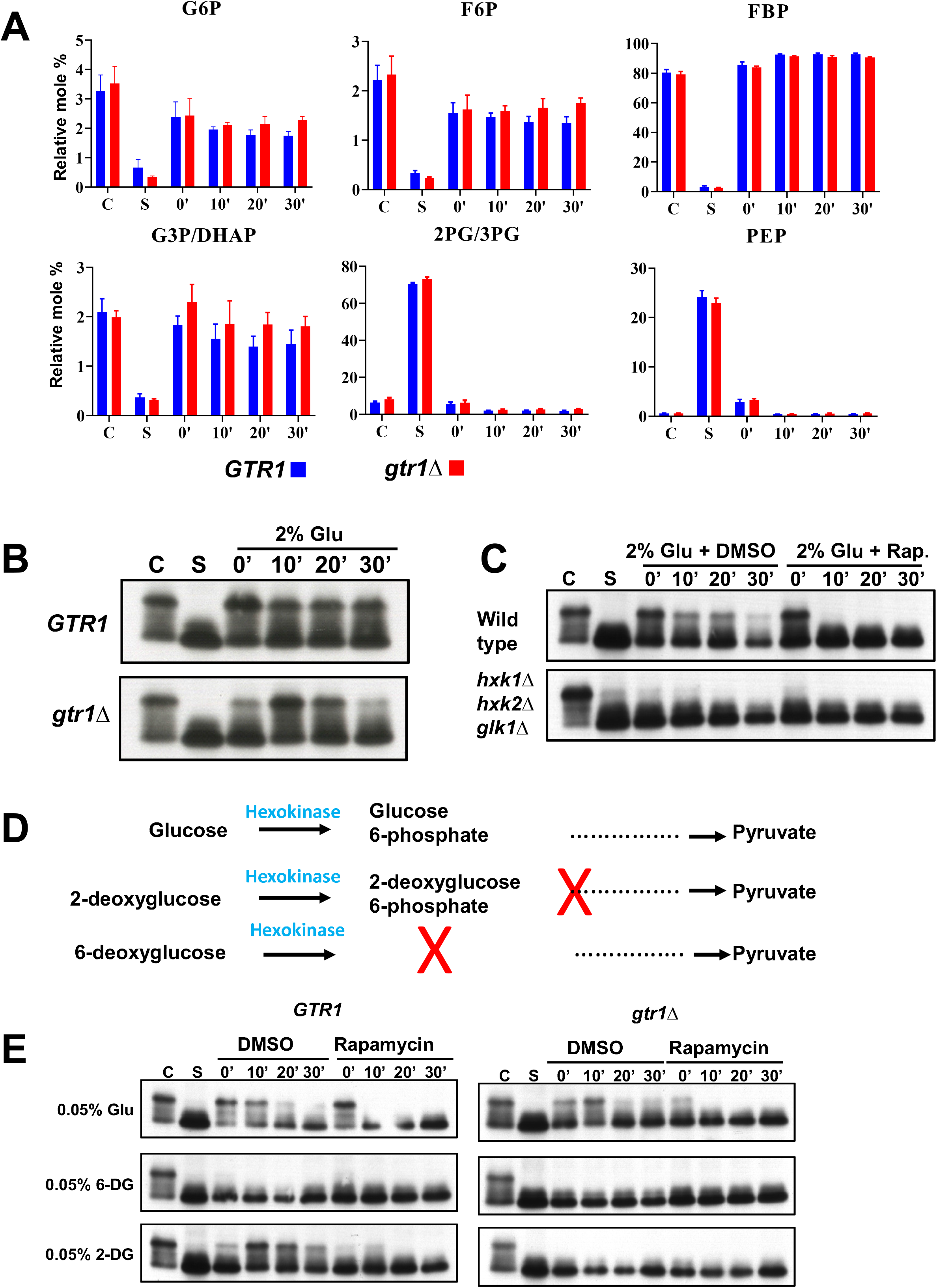
Phosphorylation of Glucose is required for glucose-induced TORC1 activation. A. Wild type and *gtr1Δ* cells were grown in SC-Glu medium in logarithmic phase (C) and then subjected to complete nutrient starvation by incubating them in water for 1 hour. Starved cells (S) were transferred to a solution containing 2% glucose. Aliquots of the cultures were taken after 0’, 10’, 20’ and 30’ for preparing protein extracts and for extracting metabolites for LC-MS analysis. Relative molar levels of various glycolytic metabolites in wild type and *gtr1Δ* cells at different stages of glucose-induced TORC1 activation assay are shown. B.Sch9 phosphorylation in wild type and *gtr1Δ* cells described in A was assayed by Western blotting. C.Wild type cells and hexokinase-deficient triple mutant cells were grown to logarithmic phase (C) in SC-Raf/Gal medium (2% raffinose and 2% galactose) and were subjected to complete nutrient starvation by incubating them in water for 1 hour. Starved cells (S) were then transferred to a solution containing 2% glucose in the presence and absence of rapamycin (2 µM). Aliquots of the cultures were taken after 0’, 10’, 20’ and 30’ and used for preparing protein extracts. Phosphorylation of Sch9 was monitored by Western blotting. D.Schematic showing the metabolic fates of glucose, 2-deoxyglucose and 6-deoxyglucose. E.Wild type cells and *gtr1Δ*cells in logarithmic phase (C) in SC-Glu medium were subjected to complete nutrient starvation by incubating them in water for 1 hour. Starved cells (S) were then transferred to a solution containing 0.05% Glucose, 0.05% 2-DG or 0.05% 6-DG in the presence and absence of rapamycin (2 µM). Aliquots of the cultures were taken after 0’, 10’, 20’ and 30’ and were used for preparing protein extracts. Phosphorylation of Sch9 was monitored by Western blotting.

To investigate whether metabolism of glucose is required for TORC1 activation, we compared the ability of glucose to activate TORC1 in wild type and hexokinase-deficient triple mutant (*hxk1Δ hxk2Δ glk1Δ*) cells. Addition of glucose resulted in rapamycin-sensitive Sch9 phosphorylation in wild type cells but not in hexokinase-deficient cells (Figure 1C) suggesting that phosphorylation of glucose to glucose 6-phosphate is essential for glucose-induced TORC1 activation. To confirm this result, we tested the ability of two glucose analogues namely 2-deoxyglucose (2-DG) and 6-deoxyglucose (6-DG) to activate TORC1. 6-DG cannot be phosphorylated by hexokinase and is therefore not metabolised (Figure 1D). On the other hand, 2-DG can be phosphorylated by hexokinase forming 2-deoxyglucose 6-phosphate but cannot be metabolised further (Figure 1D). We observed that 2% 2-DG (but not 2% 6-DG) activated TORC1 albeit weakly compared to 2% glucose (Supplementary Figure S1A). As 2-DG is a potent inhibitor of glycolysis and depletes cellular ATP levels (Kondo and Beutler, 1979), we tested lower amounts of 2-DG (ranging from 0.2% to 0.0125%) in the TORC1 activation assay. We found that 0.05% 2-DG was most effective in activating TORC1 (Supplementary Figure S1B). We compared the abilities of 0.05% of 2-DG, 6-DG and glucose to activate TORC1. Addition of 0.05% glucose or 0.05% 2-DG to starved cells resulted in increased Sch9 phosphorylation that was abolished by rapamycin treatment (Figure 1E). However, 6-DG failed to activate TORC1 (Figure 1E and Supplementary Figure S1A). These results indicate that hexokinase-mediated conversion of glucose to glucose 6-phosphate is essential for TORC1 activation.

We then compared the ability of 0.05% glucose and 2-DG to activate TORC1 in *gtr1Δ* cells. Interestingly, 0.05% glucose but not 0.05% 2-DG was able to activate TORC1 in *gtr1Δ* cells (Figure 1E). As 2-DG cannot proceed beyond the first step in glycolysis, our results imply that metabolism of glucose beyond glucose 6-phosphate is required for TORC1 activation *via* the Gtr1-independent pathway. On the other hand, conversion of glucose to glucose 6-phosphate is sufficient for Rag GTPase-dependent pathway for TORC1 activation.

Glucose 6-phosphate can be further metabolized either via the glycolytic pathway or the pentose phosphate pathway (PPP). To test which pathway is required for Gtr1-independent TORC1 activation by glucose, we deleted *ZWF1* (which encodes the glucose-6-phosphate dehydrogenase that catalyses the first committed step in the PPP) or *PFK1* (which encodes phosphofructokinase that catalyses first committed step in glycolysis). Deletion of *ZWF1* had no major effect on glucose-induced TORC1 activation in wild type and *gtr1*Δ strains (Figure S2A) indicating that the PPP is not essential for glucose-induced TORC1 activation. However, deletion of *PFK1* greatly reduced glucose-induced TORC1 activation in the wild type strain (Supplementary Figure S2B) and completely abolished TORC1 activation in the *gtr1Δ* strain (Supplementary Figure S2B). These results suggest that glycolysis is required for Rag GTPase-independent pathway for glucose-induced TORC1 activation. This is consistent with the failure of 2-DG to activate TORC1 in *gtr1*Δ cells. Although not essential, glycolysis is also required for efficient glucose-induced TORC1 activation via the Rag GTPase-dependent pathway.

### Use of SEACIT mutants uncovers two pathways of Rag GTPase-dependent TORC1 activation by glucose

To confirm the function of Gtr1 in glucose-induced TORC1 activation pathways, we tested the role of SEACIT complex in TORC1 activation. SEACIT complex is composed of Iml1/Npr2/Npr3 and inhibits TORC1 by promoting hydrolysis of GTP bound to Gtr1. We tested the ability of glucose and 2-DG to activate TORC1 in wild type and *npr3Δ* /*npr2Δ* / *iml1Δ* strains that are deficient in SEACIT function. As expected, glucose-induced TORC1 activation was stabilised in SEACIT mutants in comparison to the wild type strain (Figure 2A). Deletion of *GTR1* abolished the stabilising effect of SEACIT mutations on TORC1 activation (Supplementary Figure S3). These results support the notion that stabilization of Gtr1^GTP^ in SEACIT-deficient mutants boosts TORC1 activation. Much to our surprise, we found that 2-DG-induced activation of TORC1 was abolished in SEACIT mutant strains (Figure 2A and Supplementary Figure S3A). Although the Gtr1^GTP^ form boosts glucose-induced TORC1 activation, it is deficient in 2-DG induced TORC1 activation. To confirm this result, we constructed *gtr1Δ gtr2Δ* strains expressing either wild type Gtr1/Gtr2 or various mutant heterodimers that bind GTP or GDP (Gtr1^GTP^/ Gtr2^GDP^, Gtr1^GDP^/ Gtr2^GTP^, Gtr1^GTP^/ Gtr2^GTP^, Gtr1^GDP^/ Gtr2^GDP^). We used *gtr1Δ gtr2Δ* strains containing an empty vector as a negative control. As expected, the canonical active form Gtr1^GTP^/ Gtr2^GDP^ stabilized glucose-induced TORC1 activation in comparison to cells expressing wild type Gtr1/Gtr2 (Figure 2B). In strain expressing the inactive form Gtr1^GDP^/ Gtr2^GTP^, glucose-induced TORC1 activation was delayed as seen in *gtr1Δ gtr2Δ* cells. Consistent with our results with SEACIT mutants, 2-DG failed to activate TORC1 in the Gtr1^GTP^/Gtr2^GDP^ expressing strain (Figure 2B). Remarkably, 2-DG activated TORC1 in cells expressing Gtr1^GTP^/ Gtr2^GTP^ (Figure 2B). These results indicate that glucose activates TORC1 through Gtr1/Gtr2 *via* at least two pathways. In the first pathway, the Gtr1^GTP^ /Gtr2^GDP^ activates TORC1 upon glucose addition which we refer to this as the canonical pathway. This pathway requires metabolism beyond glucose 6-phosphate as 2-DG fails to stimulate this. In the second pathway, 2-DG activates TORC1 possibly via Gtr1^GTP^/ Gtr2^GTP^ which we refer to as the ‘non-canonical pathway’ for the remainder of this manuscript.

**Figure 2.**
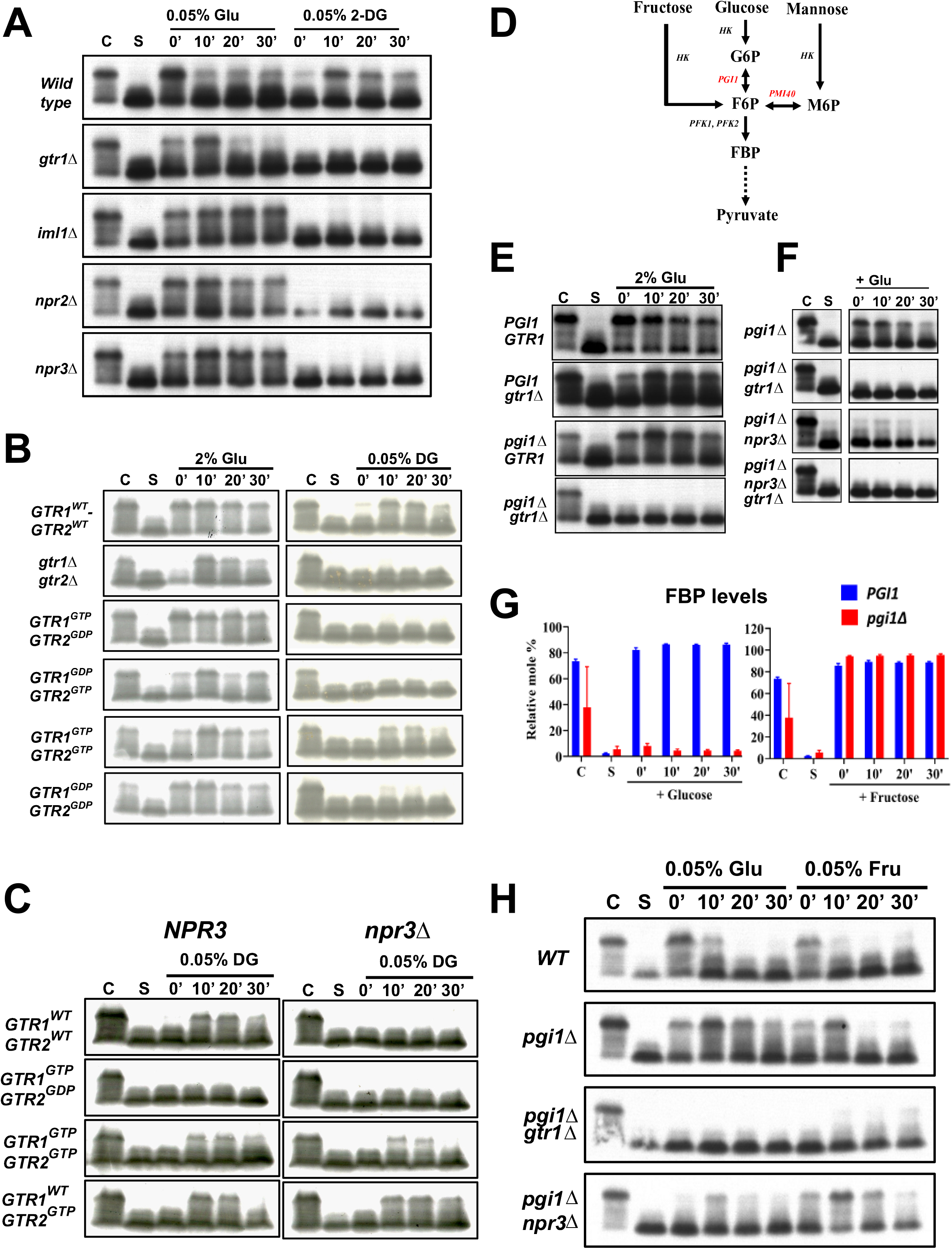
Glucose 6-phosphate activates the non-canonical pathway of TORC1 activation via Gtr1^GTP^/Gtr2^GTP^. A.Wild type, *gtr1Δ, iml1Δ, npr2Δ* and *npr3Δ* cells were grown in logarithmic phase (C) in SC-Glu medium were subjected to complete nutrient starvation by incubating them in water for 1 hour. Starved cells (S) were then transferred to a solution containing 0.05% Glucose or 0.05% 2-DG. Aliquots of the cultures were taken after 0’, 10’, 20’ and 30’ and were used for preparing protein extracts. Phosphorylation of Sch9 was monitored by Western blotting. B.*gtr1Δ gtr2Δ* cells expressing different combinations of Gtr1/ Gtr2 variants (Wild type, GTP, GDP or empty vector) from the tetracycline-inducible promoter were grown to logarithmic phase (C) in SC-Glu medium + tetracycline medium and were subjected to complete nutrient starvation by incubating them in water for 1 hour. Starved cells (S) were then transferred to a solution containing 0.05% Glucose or 0.05% 2-DG. Aliquots of the cultures were taken after 0’, 10’, 20’ and 30’ and were used for preparing protein extracts. Phosphorylation of Sch9 was monitored by Western blotting. C.Wild type and *npr3Δ* cells expressing different combinations of Gtr1/ Gtr2 variants (Gtr1^WT^-Gtr2^WT^, Gtr1^GTP^-Gtr2^GDP^, Gtr1^GTP^-Gtr2^GTP^ and Gtr1^WTP^-Gtr2^GTP^) from the tetracycline-inducible promoter were treated with either 0.05% glucose or 0.05% 2-DG in the TORC1 activation assay and analysed as described in B. D.Fructose and mannose but not glucose can enter glycolysis without phosphoglucoisomerase. E.Wild type, *gtr1Δ, pgi1Δ* and *pgi1Δ gtr1Δ* cells were grown to logarithmic phase (C) in SC-Raf/Gal medium and were subjected to complete nutrient starvation by incubating them in water for 1 hour. Starved cells (S) were then transferred to a solution containing 2% glucose. Aliquots of the cultures taken after 0’, 10’, 20’ and 30’ were used for preparing protein extracts. Phosphorylation of Sch9 was monitored by Western blotting. F.*pgi1Δ, pgi1Δ npr3Δ, pgi1Δ gtr1Δ* and *pgi1Δ npr3Δ gtr1Δ* cells were tested in the glucose-induced TORC1 activation assay as described in E. G.Wild type and *pgi1Δ* cells were grown in SC-Raf/Gal medium in logarithmic phase (C) and then subjected to complete nutrient starvation by incubating them in water for 1 hour. Starved cells (S) were transferred to a solution containing 2% glucose or 2% fructose. Aliquots of the cultures were taken after 0’, 10’, 20’ and 30’ for preparing protein extracts and for extracting metabolites for LC-MS analysis. Percentage mole fraction of FBP levels is plotted for wild type and *pgi1Δ* cells treated with 2% glucose (left panel) or 2% fructose (right panel) at the indicated time points. H.Wild type, *pgi1Δ, pgi1Δ npr3Δ* and *pgi1Δ gtr1Δ* cells were tested with 0.05% glucose and 0.05% fructose in the TORC1 activation assay as described in E.

2-DG failed to activate TORC1 in *npr3Δ* cells which is predicted to be enriched for the Gtr1^GTP^ form. This appears to inconsistent with the notion that Gtr1^GTP^/ Gtr2^GTP^ heterodimer activates TORC1 *via* the non-canonical pathway. We hypothesized that blocking hydrolysis of GTP bound to Gtr1 could either prevent GTP binding to Gtr2 or block dissociation of GDP from Gtr2. This will prevent the formation of Gtr1^GTP^/Gtr2^GTP^ heterodimers in *npr3Δ* cells and block 2-DG-induced TORC1 activation. If this hypothesis is true, then expression of the Gtr2^GTP^ mutant should suppress *npr3Δ*’s defect in 2-DG-induced TORC1 activation. We compared 2-DG-induced TORC1 activation in *npr3Δ GTR2* and *npr3Δ GTR2*^GTP^ cells (Figure 2C). 2-DG activated TORC1 in *npr3Δ* GTR2^GTP^ cells but not in *npr3Δ GTR2* cells supporting the idea that Gtr1^GTP^/Gtr2^GTP^ activates TORC1 via the non-canonical pathway (Figure 2C). Lst4 is required for hydrolysis of GTP bound to Gtr2 (Peli-Gulli et al., 2015). We found that 2-DG mediated TORC1 activation was quicker in *lst4Δ* cells compared to wild type cells further supporting the idea that Gtr1^GTP^/ Gtr2^GTP^ heterodimer activates TORC1 via the non-canonical pathway (Figure S3B).

### Glucose 6-phosphate activates TORC1 *via* a non-canonical Rag GTPase-dependent pathway

As 2-DG is a synthetic compound, the non-canonical pathway of TORC1 activation observed could be an artefact. If 2-DG induced TORC1 activation is physiological, then glucose 6-phosphate should also activate TORC1 *via* the non-canonical pathway. To test this, we constructed the phosphoglucoisomerase deletion strain (*pgi1Δ*) that cannot interconvert glucose 6-phosphate and fructose 6-phosphate (Figure 2D). Addition of glucose to *pgi1Δ* cells is expected to cause an accumulation of glucose 6-phosphate thus ‘phenocopying’ 2-DG-treated cells. Addition of glucose activated TORC1 in *pgi1Δ* cells but not in *pgi1Δ gtr1Δ* cells (Figure 2E and 2F). This is consistent with our observations that 2-DG-induced TORC1 activation is dependent on Gtr1. Crucially, addition of glucose to *pgi1Δ npr3Δ* cells resulted in drastically reduced TORC1 activation in comparison to *pgi1Δ* cells (Figure 2F). Our results support the notion that glucose 6-phosphate formation activates TORC1 *via* a Rag GTPase-dependent non-canonical pathway. Conversely, as activation by glucose (but not by 2-DG) is increased in SEACIT mutants in comparison to wild type cells, we can conclude that glucose has to be metabolized beyond glucose 6-phosphate for TORC1 activation via the Rag GTPase-dependent canonical pathway.

Fructose is phosphorylated by hexokinase to form fructose 6-phosphate and can enter glycolysis without phosphoglucoisomerase activity (Figure 2D). We confirmed this by assaying changes in levels of glycolytic metabolites caused by addition of fructose and glucose to *pgi1Δ* strains. Addition of fructose but not glucose to the *pgi1Δ* strain increased the relative molar levels of FBP (Figure 2G). By titrating the amount of fructose, we found that 0.05% fructose was optimal in activating TORC1 in the *pgi1Δ* strain (data not shown). TORC1 activation induced by 0.05% fructose was more stable in the *pgi1Δ npr3Δ* strain in comparison to the *pgi1Δ* strain (Figure 2H). In contrast, activation by 0.05% glucose was reduced in *pgi1Δ npr3Δ* cells in comparison to *pgi1Δ* cells (Figure 2H). These results support the notion that metabolism to glucose 6-phosphate activates TORC1 through a non-canonical pathway and metabolism beyond glucose 6-phosphate is required for Rag GTPase-dependent canonical pathway for TORC1 activation.

We then tested whether mannose 6-phosphate can also activate the non-canonical Rag GTPase-dependent pathway. Like fructose, mannose can enter glycolysis without phosphoglucose isomerase activity (Figure 2D). Mannose is phosphorylated by hexokinase to form mannose 6-phosphate which is then converted into fructose 6-phosphate by the phosphomannose isomerase Pmi40. We compared the kinetics of TORC1 activation induced by glucose and mannose (2% and 0.05%) in wild type, *gtr1Δ, pmi40Δ* and *pmi40Δ gtr1Δ* cells. As expected, glucose-induced TORC1 activation was comparable in wild type, *pmi40Δ* and their *gtr1Δ* variants (Supplementary Figure S4). Mannose and glucose activated TORC1 to comparable extents in wild type and *gtr1Δ* cells. Interestingly,0.05% mannose but not 2% mannose activated TORC1 in *pmi40Δ* cells. TORC1 activation caused by 0.05% mannose was abolished in *pmi40Δ gtr1Δ* cells (Supplementary Figure S4). TORC1 activation by mannose in *pmi40Δ* cells is reminiscent of TORC1 activation results obtained in wild type cells treated with 2-DG. Taken together our results indicate that formation of glucose 6-phosphate /mannose 6-phosphate is necessary and sufficient to activate the Rag GTPase-dependent non-canonical pathway.

### Formation of FBP is sufficient for the canonical Rag GTPase-dependent pathway of TORC1 activation

We then sought to identify glycolytic steps downstream of glucose 6-phosphate that is required for the Rag GTPase-dependent canonical pathway and the Rag GTPase-independent pathway of TORC1 activation. Aldolase (Fba1) catalyses the conversion of FBP into glyceraldehyde 3-phosphate (G3P) and dihydroxyacetone phosphate (DHAP). Since *FBA1* is an essential gene, we constructed a yeast strain that expresses *FBA1* from the tetracycline-repressible promoter. *PTetoff-FBA1* cells were dead on tetracycline-containing SC-Glucose medium plates (data not shown). To deplete aldolase, *PTetoff-FBA1* cells were grown overnight in synthetic medium in the presence of tetracycline. Cells from the overnight culture were inoculated at OD600 nm= 0.2 and cultured further for 5-6 hours in SC-Glucose medium + tetracycline. Strain expressing *FBA1* from the native promoter served as a control. Growth of *PTetoff-FBA1* cells was severely inhibited in the presence of tetracycline indicating an efficient depletion. We compared the relative proportions of glycolytic metabolites in wild type and *FBA1-*depleted cells following addition of glucose. Relative molar levels of FBP following glucose addition increased to 92% in wild type cells but increased to 97% in the *PTetoff-FBA1* cells (Figure 3A). Crucially, the downstream glycolytic metabolite (G3P/DHAP) was not detectable in *PTetoff-FBA1* cells indicating an efficient depletion of *FBA1* (Figure 3A). TORC1 was activated by glucose albeit at reduced levels in *PTetoff-FBA1* cells in comparison to wild type cells (Figure 3B). In contrast, TORC1 activation was completely blocked in *PTetoff-FBA1 gtr1Δ* cells (Figure 3B) indicating that metabolism of glucose beyond FBP is required for TORC1 activation *via* the Rag GTPase-independent pathway. However, formation of FBP is sufficient for Rag GTPase-dependent pathway of TORC1 activation. To confirm that the activation of TORC1 by glucose in *PTetoff-FBA1* strain was *via* the canonical pathway, we compared glucose-induced TORC1 activation in *PTetoff-FBA1* and *PTetoff-FBA1 npr3Δ* cells. TORC1 was activated following addition of glucose and remained stable in *PTetoff-FBA1 npr3Δ* cells in comparison to *PTetoff-FBA1* cells (Figure 3B). Taken together, our results suggest that formation of FBP is sufficient for TORC1 activation *via* the canonical pathway but further glycolytic steps beyond FBP formation is required for the Rag GTPase-independent pathway of TORC1 activation.

**Figure 3.**
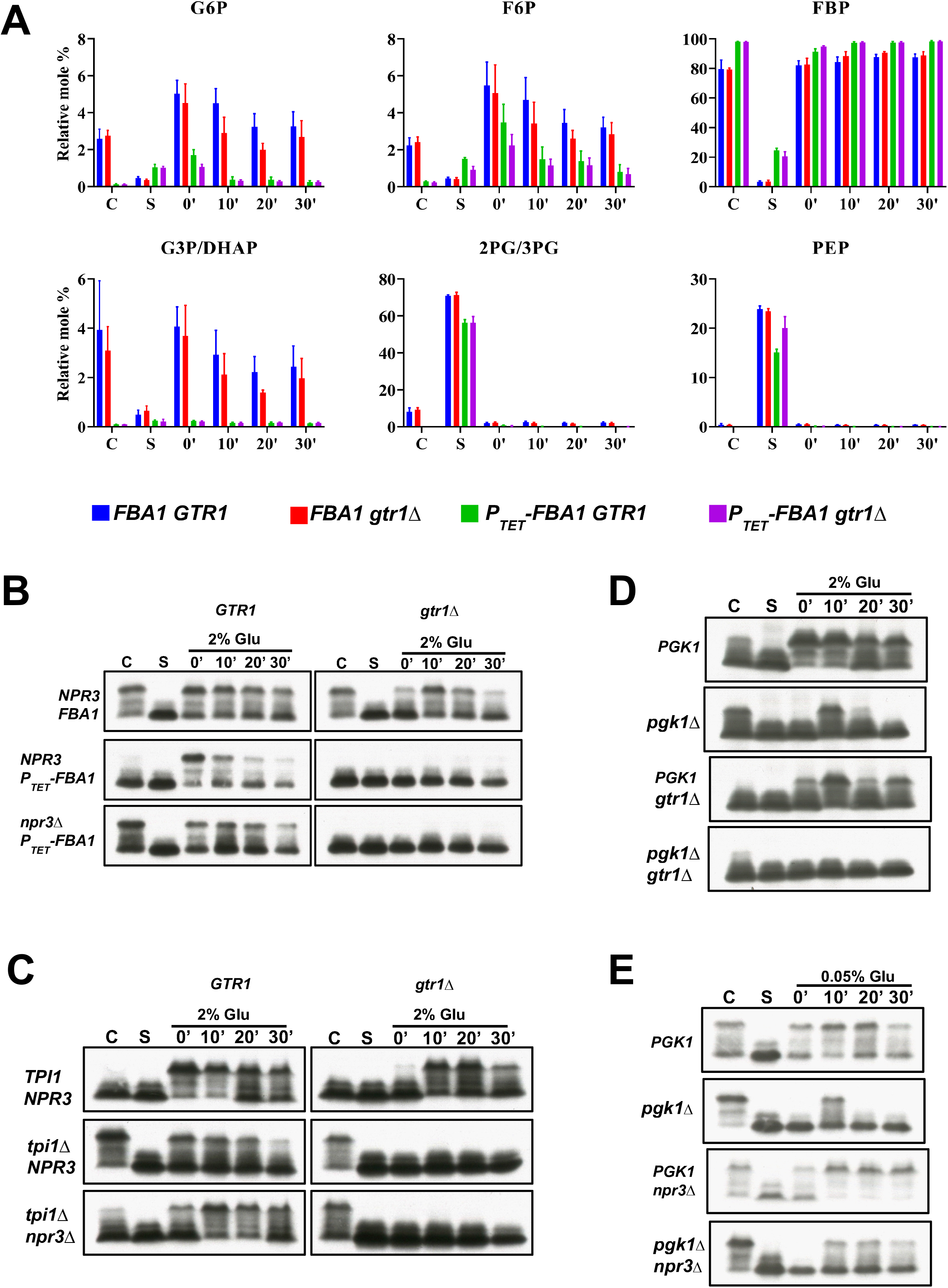
Formation of FBP from glucose is sufficient for the canonical pathway of TORC1 activation. A.Wild type and *PTetoff-FBA1* cells were grown in SC-Glu medium in the presence of tetracycline and then subjected to complete nutrient starvation by incubating them in water for 1 hour. Starved cells (S) were transferred to a solution containing 2% glucose. Aliquots of the cultures were taken after 0’, 10’, 20’ and 30’ for extracting metabolites for LC-MS analysis and for preparing protein extracts. Relative molar levels of various glycolytic metabolites in wild type and *PTetoff-FBA1* cells at different stages of glucose-induced TORC1 activation assay are shown. B.Sch9 phosphorylation in wild type and *PTetoff-FBA1* cells was assayed by Western blotting. C.Wild type and *tpi1Δ* cells were grown in SC-EtOH/Gly medium (2% ethanol and 2% glycerol) and then subjected to complete nutrient starvation by incubating them in water for 1 hour. Starved cells (S) were transferred to a solution containing 2% glucose. Aliquots of the cultures were taken after 0’, 10’, 20’ and 30’ for analysis for Sch9 phosphorylation and for extracting metabolites for LC-MS analysis. Sch9 phosphorylation in wild type and *tpi1Δ* cells was assayed by Western blotting. Relative molar levels of various glycolytic metabolites in wild type and *tpi1Δ* cells at different stages of glucose-induced TORC1 activation assay are shown in Figure S5. D.Wild type and *pgk1Δ* cells were treated as in C and Sch9 phosphorylation was assayed by Western blotting. Relative molar levels of various glycolytic metabolites in wild type and *pgk1Δ* cells at different stages of glucose-induced TORC1 activation assay are shown in Figure S6. E.Wild type, *pgk1Δ, npr3Δ* and *pgk1Δ npr3Δ* cells were analysed in the TORC1 activation assay as described in D. Sch9 phosphorylation was assayed by Western blotting.

### Complete glycolysis is essential for TORC1 activation via the Rag GTPase-independent pathway

To determine glycolytic steps downstream of FBP formation that is required for Rag GTPase-independent pathway of TORC1 activation, we first tested the role of Triosephosphate isomerase (Tpi1) which interconverts Dihydroxyacetone phosphate (DHAP) and Glyceraldehyde 3-phosphate (G3P). In *tpi1Δ* cells, G3P but not DHAP generated by fructose1,6-bisphosphate aldolase will be metabolised further in glycolysis. We analysed the relative levels of glycolytic metabolites in wild type and *tpi1Δ* cells during the glucose-induced TORC1 activation assay. As expected, levels of DHAP were high in *tpi1Δ* cells (cycling and starved) in comparison to wild type cells (Supplementary Figure S5). Addition of glucose to starved *tpi1Δ* cells caused a rise in relative FBP levels albeit to a lower extent than in wild type cells. TORC1 activation was reduced in *tpi1Δ* cells in comparison to the wild type cells (Figure 3C). However, TORC1 activation was completely abolished in *tpi1Δ gtr1Δ* cells (Figure 3C). These results suggest that net ATP production from glycolysis is required for the Rag GTPase-independent pathway of TORC1 activation. To confirm that activation of TORC1 by glucose in *tpi1Δ* cells is *via* the Rag GTPase-dependent canonical pathway, we compared TORC1 activation in *tpi1Δ* and *tpi1Δ npr3Δ* cells (Figure 3C). Glucose-induced TORC1 activation was more stable in *tpi1Δ npr3Δ* cells in comparison to *tpi1Δ* cells indicating that triosephosphate isomerase activity is not required for TORC1 activation by glucose *via* the Rag GTPase-dependent canonical pathway. These results further support the notion that formation of FBP is sufficient for activation of TORC1 *via* the canonical Rag GTPase-dependent pathway.

We then examined the ability of *pgk1Δ, gpm1Δ* and *cdc19Δ* mutants that are defective in the remaining glycolytic steps to metabolize glucose under our TORC1 assay conditions. The *cdc19Δ* and *gpm1Δ* mutant cells could not initiate glycolysis (Figure S6A). Addition of glucose to *cdc19Δ* and *gpm1Δ* cells failed to activate TORC1 consistent with the idea that glycolysis is essential for glucose-induced TORC1 activation (Figure S6B). Interestingly, we observed an accumulation of FBP in starved *pgk1Δ* cells and a further increase in FBP levels upon glucose addition (Figure S8A). Glucose (2%) activated TORC1 in wild type and *pgk1Δ* cells (Figure 3D). However, glucose failed to activate TORC1 in *pgk1Δ gtr1Δ* cells (Figure 3D). To test whether the canonical pathway is active in *pgk1Δ* cells, we compared glucose-induced TORC1 activation in *pgk1Δ* and *pgk1Δ npr3Δ* cells. We used lower concentrations of glucose (0.05%) as we reasoned that *pgk1Δ* cells might have lower amounts of ATP. TORC1 activation was stabilized in *pgk1Δ npr3Δ* cells in comparison to *pgk1Δ* cells indicating that phosphoglycerate kinase activity is not required for the canonical pathway for glucose-induced TORC1 activation (Figure 3E). These results further support the idea that conversion of glucose to FBP is sufficient for TORC1 activation *via* the canonical Rag GTPase-dependent pathway, but complete glycolysis of glucose is required for the Rag GTPase-independent pathway. It is interesting to note that starved *pgk1Δ* cells contain fructose 1,6-bisphosphate but lack TORC1 activity. This indicates that presence of FBP is not sufficient for TORC1 activation.

### Mitochondrial function is essential for the non-canonical pathway of TORC1 activation

Following glycolysis, pyruvate is converted into acetyl-CoA which generates ATP *via* the mitochondrial oxidative phosphorylation pathway. We tested whether mitochondrial function is required for glucose-induced TORC1 activation. We generated the *pet100*Δ strain that lacks mitochondrial electron transport chain function, and petite strains by ethidium bromide treatment of wild type cells. Both the *pet100Δ* and petite strains were unable to grow in medium containing a non-fermentable carbon source (data not shown) consistent with a defect in mitochondrial function. We compared the abilities of 2-DG and glucose to activate TORC1 in wild type, *pet100Δ* and their *gtr1Δ* variants. Glucose-induced TORC1 activation in *pet100Δ* and *pet100Δ gtr1Δ* strains were comparable to the wild type counterparts (Figure 4A). However, 2-DG failed to activate TORC1 in *pet100Δ* strains (Figure 4A). Glucose-induced TORC1 activation in petite strains was weaker compared to wild type strain (Figure 4B) but the relative timings of TORC1 activation in wild type and *gtr1*Δ petite strains was comparable to their wild type counterparts. However, 2-DG failed to activate TORC1 in the petite strains (Figure 4B). These results indicate that a functional mitochondrion is essential for the non-canonical pathway but not required for canonical Rag GTPase-dependent and Rag GTPase-independent pathways. To confirm this, we tested the effect of antimycin A, an inhibitor of mitochondrial respiratory chain (Slater, 1973), on the non-canonical pathway of TORC1 activation. We compared the ability of glucose and 2-DG to activate TORC1 in wild type and *pgi1Δ* cells in the presence or absence of antimycin A treatment. Glucose-induced activation in wild type and *gtr1Δ* cells was largely unaffected by antimycin A treatment (Figure 4C). However, 2-DG-induced TORC1 activation was abolished by antimycin A treatment. Likewise, activation of TORC1 by glucose in *pgi1Δ* cells (which phenocopies 2-DG treatment) was also abolished by antimycin treatment A (Figure 4C). These results demonstrate that active mitochondrial function is essential for the non-canonical pathway of TORC1 activation but dispensable for the Rag GTPase-dependent canonical/ and Rag GTPase-independent pathways of TORC1 activation.

**Figure 4.**
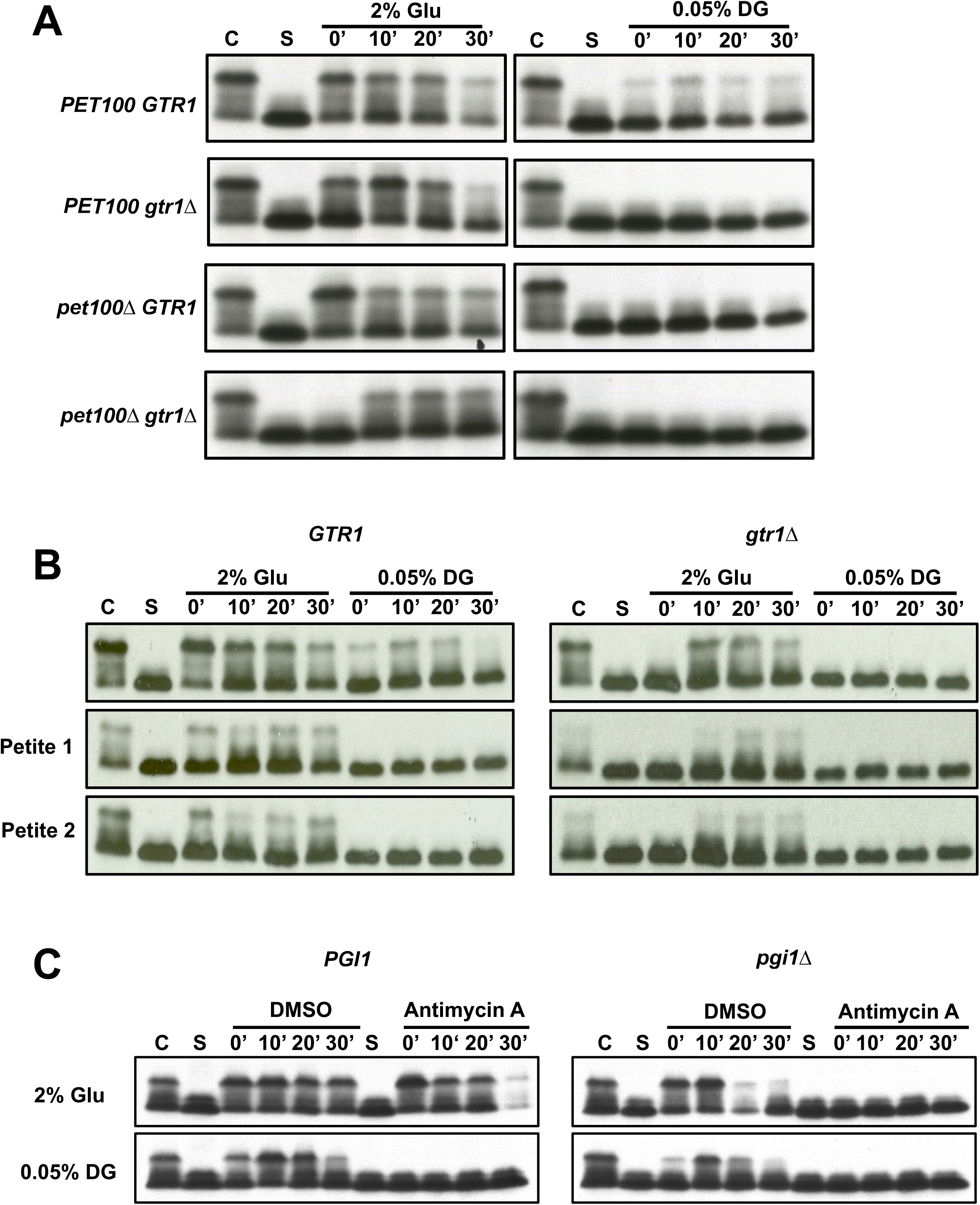
Mitochondrial function is required for the non-canonical pathway of glucose-induced TORC1 activation. A.Wild type, *gtr1Δ, pet100Δ* and *pet100Δ gtr1Δ* cells grown to logarithmic phase (C) in SC-Glu medium were subjected to complete nutrient starvation by incubating them in water for 1 hour. Starved cells (S) were then transferred to a solution containing 2% Glucose or 0.05% 2-DG. Aliquots of the cultures were taken after 0’, 10’, 20’ and 30’ and were used for preparing protein extracts. Phosphorylation of Sch9 was monitored by Western blotting. B.Same as described in A but performed with wild type, *gtr1Δ* and two petite strains with or without *gtr1Δ*. C.Same as described in A but performed with wild type, *gtr1Δ, pgi1Δ* and *pgi1Δ gtr1Δ* cells with or without antimycin treatment (50 µM).

### Known FBP-binding proteins are not required for the canonical pathway of TORC1 activation

FBP formation is essential for glucose-induced TORC1 activation *via* the canonical pathway. FBP could activate TORC1 by directly binding to its subunits or indirectly *via* proteins that regulate TORC1 activity. Alternatively, generation of FBP from glucose could modify the cellular milieu which then activates TORC1. We explored the role of three FBP-binding proteins namely the aldolase, pyruvate kinase and fructose 1,6-bisphosphatase in glucose-induced TORC1 activation. FBP binds to aldolase and inactivates AMPK in human cells (Zhang et al., 2017). To test if the yeast aldolase has a non-enzymatic role in glucose-induced TORC1 activation, we replaced the yeast aldolase with the human aldolase. Although yeast and human aldolases catalyse the same enzymatic reaction, they are not evolutionarily related and have no detectable sequence homology (Marsh and Lebherz, 1992). Human aldolase rescued the lethal phenotype of yeast *fba1* deletion (data not shown). Glucose induced TORC1 activation in the human-aldolase expressing yeast strain and was comparable to activation in wild type yeast cells (Supplementary Figure S7A). Furthermore, TORC1 activation was enhanced in *npr3Δ FBA1*^*huma*n^ cells (Supplementary Figure S7A). These results strongly suggest that aldolase does not have a non-catalytic role in TORC1 activation in yeast. Deletion of *FBP1* gene that encodes the fructose1,6-bisphosphatase did not affect glucose induced TORC1 activation (Supplementary Figure S7B). Moreover, *npr3*Δ stabilized glucose-induced TORC1 activation in *fbp1Δ* cells (Supplementary Figure S7B) indicating that activation of the canonical pathway does not require fructose 1,6-bisphosphatase.

Cdc19’s enzymatic activity is induced upon FBP binding (Supplementary Figure S7C). To test if FBP binding to Cdc19 is required for glucose-induced TORC1 activation, we constructed a catalytically active variant of Cdc19 that is deficient in FBP binding. R459 in Cdc19 forms an electrostatic interaction with the 1’-phosphate group of FBP (Jurica et al., 1998). R459Q mutation in Cdc19 abolishes FBP binding (Supplementary Figure S7C) (Fenton and Blair, 2002). Strain expressing Cdc19^R459Q^ was unable to grow in nutrient medium with glucose as a carbon source indicating that R459Q inhibits Cdc19’s enzymatic activity. E392A mutation in Cdc19 renders the catalytic activity independent of FBP binding (Fenton and Blair, 2002; Xu et al., 2012). We tested whether the E392A mutation could restore the enzymatic activity of the Cdc19^R459Q^ mutant. Indeed, a strain expressing Cdc19^E392A R459Q^ mutant was able to grow in glucose-containing medium. TORC1 activation induced by glucose in Cdc19^WT^ and Cdc19^E392AR459Q^ strains were comparable (Supplementary Figure S7C). These results strongly suggest that the Cdc19’s ability to bind FBP is not required for glucose-induced TORC1 activation.

### Vacuolar ATPase activity is required for Rag GTPase-independent pathway of TORC1 activation

Vacuolar ATPase is a multi-subunit complex that uses energy derived from ATP hydrolysis to pump protons into the vacuole and maintains the low pH inside the yeast vacuoles (Hayek et al., 2019). Vacuolar ATPase disassembles during glucose starvation and reassembles upon glucose re-addition in yeast (Hayek et al., 2019). Vacuolar ATPase reassembly in yeast is facilitated by the RAVE (regulator of the ATPase of vacuolar and endosomal membranes) complex composed of Rav1, Rav2 and Skp1 (Smardon et al., 2014; Smardon et al., 2002). To test whether vacuolar ATPase reassembly is required for glucose induced TORC1 activation, we compared TORC1 activation in wild type, *gtr1Δ, rav2Δ* and *rav2Δ gtr1Δ* cells. While glucose induced TORC1 activation in wild type and *rav2Δ* to comparable extents (Figure 5A), the *rav2Δ gtr1Δ* cells were severely compromised in glucose-induced TORC1 activation (Figure 5A). These results suggest that vacuolar ATPase reassembly is required for Rag-GTPase-independent TORC1 activation pathway (Figure 5A).

**Figure 5.**
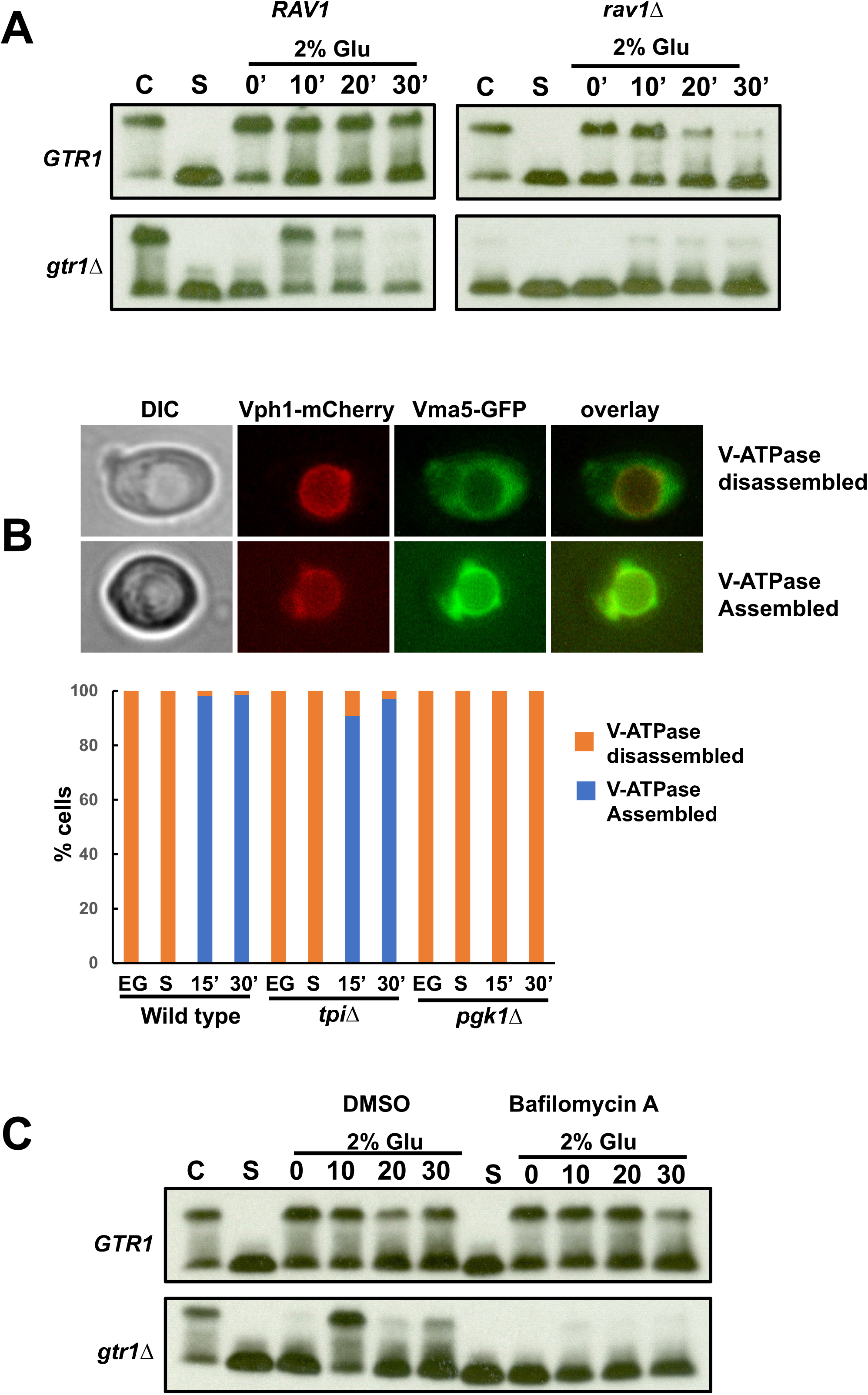
Vacuolar ATPase reassembly and activation is required for the Rag GTPase-independent pathway of TORC1 activation. A.Wild type, *gtr1Δ, rav2Δ* and *rav2Δ gtr1Δ* cells were grown in logarithmic phase (C) in SC-Glu medium were subjected to complete nutrient starvation by incubating them in water for 1 hour. Starved cells (S) were then transferred to a solution containing 2% glucose. Aliquots of the cultures were taken after 0’, 10’, 20’ and 30’ and were used for preparing protein extracts. Phosphorylation of Sch9 was monitored by Western blotting. B.Wild type, *tpi1Δ* and *pgk1Δ* cells expressing Vma5-mCherry and Vph1-GFP were subjected to the glucose-induced TORC1 activation assay. Aliquots of the cultures were taken at different points and examined by fluorescence microscopy. Percentage cells showing localization of Vma5-mCherry and Vph1-GFP were plotted. Representative images showing cells with assembled and dissembled V-ATPase are shown. C.Same as described in A but performed with wild type and *gtr1*Δ cells in the presence and absence of bafilomycin (10 µM).

To test whether vacuolar ATPase reassembles during the glucose-induced TORC1 activation assay, we tagged the cytosolic (Vph1) and membrane (Vma5) subunits of the V-ATPase with mCherry and GFP, respectively. We tested the effect of two glycolytic mutations *tpi1Δ* and *pgk1Δ* on glucose-induced vacuolar ATPase reassembly. We grew the wild type, *tpi1Δ* and *pgk1Δ* cells in ethanol-glycerol medium to log phase and then subjected them to complete nutrient starvation for 1 hour. We then added glucose (2%) to starved cells and assayed vacuolar ATPase reassembly 15 and 30 minutes after glucose addition. Vacuolar ATPase was disassembled in cells growing in ethanol-glycerol medium (Figure 5B). Vacuolar ATPase assembly following glucose addition occurred in both wild type and *tpi1Δ* cells but not in *pgk1Δ* cells. This suggests that complete glycolysis but not ATP generation per se is required for glucose-induced vacuolar ATPase assembly. However, glucose-induced TORC1 activation via the Rag GTPase-independent pathway was defective in both *tpi1Δ* (Figure 5B) and *pgk1Δ* (Figure 5B) cells. Taken together, these results suggest that complete glycolysis with net ATP production is essential for TORC1 activation via the Rag GTPase-independent pathway.

ATP generated by glycolysis could be required for V-ATPase activity. To test whether V-ATPase activity is required for glucose-induced TORC1 activation, we treated nutrient-starved cells with vacuolar ATPase inhibitor bafilomycin A1(Bowman et al., 1988) and added glucose to wild type and *gtr1Δ* cells and assayed TORC1 activation. Bafilomycin A1 treatment had little or no effect of TORC1 activation in wild type cells (Figure 5C). However, glucose-induced TORC1 activation was severely reduced in bafilomycin A1-treated *gtr1Δ* cells (Figure 5C). Taken together, our results suggest that vacuolar ATPase assembly and activation following glucose addition is required for the Rag GTPase-independent pathway for TORC1 activation.

### AMPK regulates TORC1 activation independently of Kog1 phosphorylation

AMP-activated protein kinase (AMPK) is a conserved energy sensor in eukaryotic cells (Conrad et al., 2014; Herzig and Shaw, 2018). AMPK in yeast is a heterotrimeric complex composed of the catalytic α subunit Snf1, a regulatory β subunit (Gal83/Sip1/Sip2) and the activating γ subunit Snf4 (Conrad et al., 2014). AMPK has been reported to antagonize TORC1 by phosphorylating Kog1 during glucose starvation in yeast (Hughes Hallett et al., 2015). During glucose starvation, Snf1 was reported to trigger TORC1 disassembly and relocation of TORC1 subunit Kog1/Raptor to a single body called as Kog1 body that borders the vacuole (Hughes Hallett et al., 2015). Snf1-mediated phosphorylation of Kog1 at Ser491/494 was proposed to drive this phenomenon (Hughes Hallett et al., 2015). We compared the ability of cells expressing wild type Kog1 and Kog1-S491A, S494A to both glucose and 2-DG. TORC1 activation induced by glucose was comparable in wild type and Kog1-S491A,S494A strains (Figure 6A). These results indicate that Snf1 regulates glucose-induced TORC1 activation independently of Kog1 phosphorylation.

**Figure 6.**
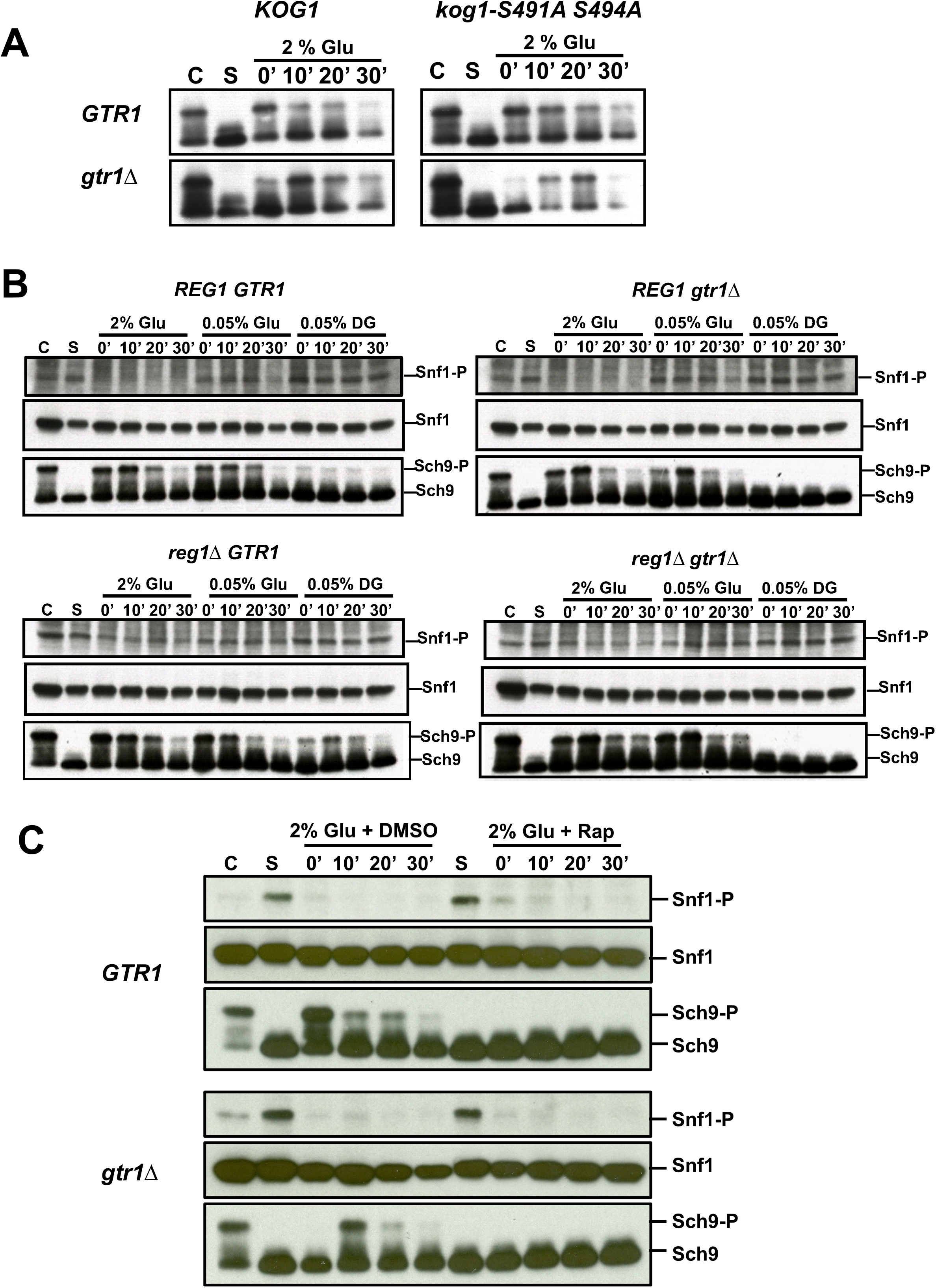
AMPK inactivation and TORC1 activation following glucose addition are not mechanistically linked. A.Wild type, *gtr1Δ, kog1-S491AS494A* and *kog1-S491AS494A gtr1Δ* cells were grown to logarithmic phase (C) in SC-Glu medium and were subjected to complete nutrient starvation by incubating them in water for 1 hour. Starved cells (S) were then transferred to a solution containing 2% glucose. Aliquots of the cultures taken after 0’, 10’, 20’ and 30’ were used for preparing protein extracts. Phosphorylation of Sch9 was monitored by Western blotting. B.Wild type, *gtr1Δ, reg1Δ* and *reg1Δ gtr1Δ* cells expressing Snf1-PK9 were subjected to the TORC1 activation assay. Aliquots of the cultures were taken after 0’, 10’, 20’ and 30’ and were used for preparing protein extracts. Snf1 phosphorylation, total Snf1 levels and Sch9 phosphorylation was monitored by Western blotting. C.Wild type and *gtr1Δ* cells grown in logarithmic phase (C) in SC-Glu medium were subjected to complete nutrient starvation by incubating them in water. After half an hour of starvation either DMSO or Rapamycin (2µM) were added to the starved cells. After another 30’ DMSO/rapamycin starved cells (S) were transferred to a solution containing 2% glucose containing either DMSO or rapamycin. Aliquots of the cultures were taken after 0’, 10’, 20’ and 30’ and were used for preparing protein extracts. Snf1 phosphorylation, total Snf1 levels and Sch9 phosphorylation was monitored by Western blotting.

### AMPK inactivation and TORC1 activation following glucose re-addition are independent events

To explore any interplay in the regulation of TORC1 and AMPK activities, we monitored AMPK activity in wild type, *gtr1Δ, reg1Δ* and *reg1Δ gtr1Δ* cells during the course of glucose-induced TORC1 activation assay. We tagged Snf1 with Pk epitope to measure the total Snf1 levels in cells by western blotting. To assay AMPK activation, we used the phospho-specific Snf1 antibody. In both wild type and *gtr1*Δ strains, Snf1 was inactive in log phase cells but was activated during complete nutrient starvation (Figure 6B). This is consistent with the well-established antagonistic role of glucose in AMPK activation. As expected, TORC1 was activated by 2% glucose in wild type cells, and in *gtr1Δ* cells albeit with a 10-minute delay (Figure 6A). In contrast, addition of 2% glucose resulted in immediate Snf1 inactivation in both wild type and *gtr1Δ* cells (Figure 6B). Interestingly, addition of either 0.05% glucose or 0.05% 2-DG to starved cells failed to inactivate Snf1. However, 0.05% glucose activated TORC1 in both wild type and *gtr1*Δ cells and 0.05% DG activated TORC1 only in wild type but not in *gtr1*Δ cells (Figure 6B). These results indicate the differing requirements of glucose for AMPK inactivation and TORC1 activation in our assay thereby uncoupling TORC1 activation from AMPK inactivation. Moreover, AMPK was constitutively active in in *reg1Δ* and *reg1Δ gtr1Δ* cells regardless of the presence or absence of glucose. However, TORC1 activation in *reg1*Δ / *reg1*Δ *gtr1*Δ cells and their corresponding wild type counterparts was comparable. Taken together, our results indicate that activation of TORC1 and inactivation of AMPK following glucose addition to starved cells are not causally linked events.

### AMPK inactivation following glucose re-addition is independent of TORC1 activity

While AMPK inactivation is not required for TORC1 activation, it is possible that TORC1 activation is required for AMPK inactivation. To test this, we treated starving wild type and *gtr1*Δ cells with either DMSO or rapamycin for 30’ to inhibit TORC1 before glucose addition. Following starvation, we added 2% glucose along with DMSO or rapamycin to the corresponding samples. Following addition of 2% glucose, TORC1 was activated in both DMSO-treated wild type and *gtr1Δ* cells (with a delay) but completely inhibited in rapamycin-treated wild type/ *gtr1Δ* cells (Figure 6C). However, addition of 2% glucose inactivated AMPK in both rapamycin- and DMSO-treated wild type/ *gtr1Δ* cells (Figure 6C). These results indicate that TORC1 activation is not required for AMPK inactivation following glucose addition to starved yeast cells.

## Discussion

An exciting yet formidable challenge in biology is to understand how a cell senses the presence of nutrients and fine tunes its growth and developmental state. Discovery and characterization of TORC1 was a significant step towards meeting this challenge in eukaryotic cells. TORC1, a highly conserved eukaryotic protein complex integrates sensory inputs from growth factors, amino acids and glucose with rates of cellular growth, metabolism and proliferation. But the molecular details of how nutrients modulate TORC1 activity remain to be uncovered. Discovery of sestrins provided a mechanistic basis for regulation of mTORC1 activity by amino acids in mammalian cells. However, how amino acids and glucose regulate TORC1 activity in *Saccharomyces cerevisiae* remain largely unknown. In this paper, we combined genetics with targeted metabolite analysis to identify cellular factors and metabolic requirements for glucose-induced TORC1 activation in *Saccharomyces cerevisiae*. We show that metabolism of glucose via the glycolytic pathway activates TORC1 through three distinct pathways (Figure 7).

**Figure 7.**
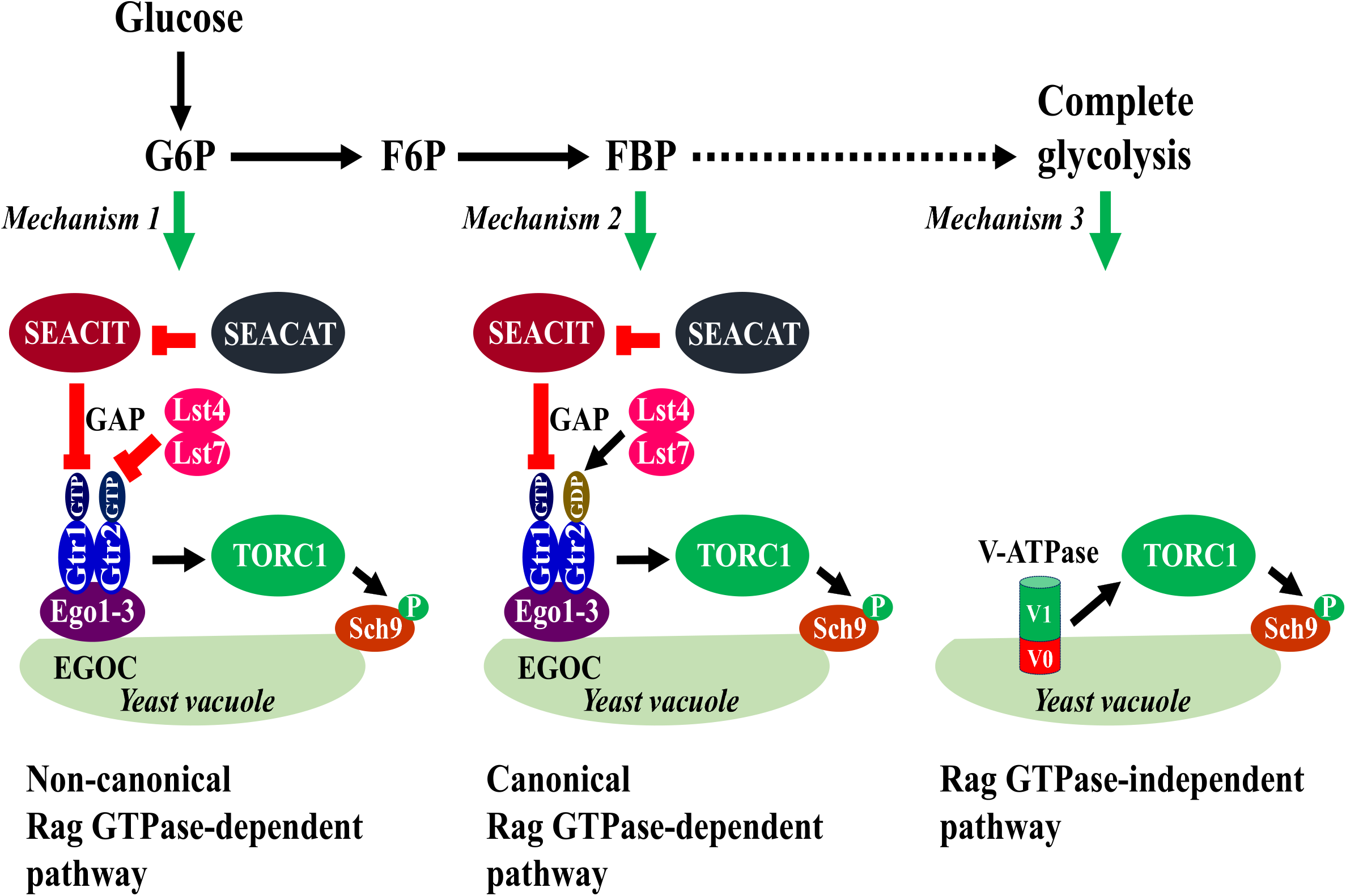
A model for glucose-induced TORC1 activation.

### A non-canonical pathway for TORC1 activation

Activation of TORC1 via binding of Gtr1^GTP^/Gtr2^GDP^ under conditions of amino acid sufficiency constitutes the canonical pathway of TORC1 activation, which is conserved from yeast to humans. Our work has uncovered a hitherto unreported non-canonical pathway of TORC1 activation that is promoted by Gtr1^GTP^/Gtr2^GTP^ but inhibited by Gtr1^GTP^/Gtr2^GDP^. We have multiple lines of evidence in support for the non-canonical pathway. Either inactivation of SEACIT complexes or expression of Gtr1^GTP^/Gtr2^GDP^ blocked of TORC1 activation in 2-DG-treated wild type cells. Addition of glucose to *pgi1Δ* cells phenocopied the 2-DG tested wild type cells. The non-canonical pathway was active in 2-DG treated cells expressing either wild type Gtr1/Gtr2 or Gtr1^GTP^/Gtr2^GTP^. SEACIT mutant cells accumulate Gtr1^GTP^ but failed to activate the non-canonical pathway. We reasoned that hydrolysis of GTP bound to Gtr1 might help GTP binding/GDP dissociation from Gtr2 thereby preventing the formation of Gtr1^GTP^/Gtr2^GTP^ in SEACIT mutant cells. Consistent with this possibility, the defect of SEACIT mutant cells to activate the non-canonical pathway was suppressed by expression of Gtr2^GTP^ bound form. As SEACIT mutant cells accumulate Gtr1^GTP^ bound form, this further shows that Gtr1^GTP^/Gtr2^GTP^ promotes the non-canonical pathway of TORC1 activation.

What might be the function of non-canonical pathway? One possibility is that yeast cells might use two different pathways to activate TORC1 during glycolysis and gluconeogenesis. During glycolysis, F16BP activates TORC1 *via* the canonical pathway. During gluconeogenesis, G6P might activate TORC1 *via* the non-canonical pathway. Interestingly, mitochondrial function essential for energy generation during gluconeogenesis is required for the non-canonical pathway. Having two separate mechanisms for TORC1 activation could facilitate specificity of downstream responses during glycolysis and gluconeogenesis. Uncovering the biological relevance of the non-canonical pathway of TORC1 activation and whether this pathway is conserved in mammalian cells are exciting challenges for the future. Intriguingly, glucose 6-phosphate activates mTORC1 in mammalian cells (Roberts et al., 2014). As observed in yeast, mTORC1 was activated by lower concentrations of 2-DG but inhibited at higher concentrations (Roberts et al., 2014). Upon G6P deprivation, hexokinase-II bound and inhibited TORC1 through its TOS motif. In the presence of glucose, G6P is formed which binds to HK-II and releases it from TORC1 thus relieving the inhibition. Testing whether 2-DG activation in mammalian cells is dependent on canonical/non-canonical states of RagA/B-RagC/D heterodimer would be informative.

How might the non-canonical pathway work? One possibility is that the TORC1 in starved cells adopts a different higher order structure which requires Gtr1^GTP^-Gtr2^GTP^ instead of Gtr1^GTP^-Gtr2^GDP^ for activation. Glucose 6-phosphate could either directly or indirectly alter TORC1 complex structure to facilitate interaction with the Gtr1-GTP:Gtr2-GTP. TORC1 forms TOROIDs during starvation in yeast which requires the inactive heterodimer namely Gtr1^GDP^-Gtr2^GTP^. However, we did not observe any TORC1 foci during complete nutrient starvation (Mohammad Alfatah and Prakash Arumugam, unpublished observations). Also, expression of Gtr1^GDP^-Gtr2^GTP^ did not activate the non-canonical pathway (Figure 2B).

### Canonical pathway of glucose-induced TORC1 activation

Activation of canonical pathway requires active Gtr1^GTP^:Gtr2^GDP^ and formation of fructose 1,6-bisphosphate. Inactivation of the SEACIT complex or expression of Gtr1^GTP^:Gtr2^GDP^ boosted the canonical pathway. While blocking glycolysis before FBP formation prevented the activation of this pathway, blocking glycolysis downstream of FBP formation did not affect this pathway. Inactivation of aldolase which boosted FBP levels hyper-activated TORC1 although transiently.

How might FBP activate TORC1? FBP was shown to inactivate AMPK in human cells by binding to aldolase (Zhang et al., 2017). It is very unlikely that FBP binds to aldolase in yeast to activate TORC1 *via* a non-enzymatic function. Although yeast aldolase and human aldolase A catalyse the same enzymatic reaction, they are not homologues and evolved convergently to perform the same function. Human aldolase was able to substitute for yeast aldolase functionally in both glycolysis and TORC1 activation. FBP also binds with other two proteins namely Fbp1 and Cdc19 which have FBPase and pyruvate kinase activity, respectively. Deletion of *FBP1* which encodes FBPase did not affect TORC1 activation. FBP binds to Cdc19 and activates its enzymatic activity allosterically. However, a constitutively active form of Cdc19 that lacks FBP binding was comparable to wild type Cdc19 in terms of activating the canonical pathway of glucose-induced TORC1 activation. *pgk1Δ* cells accumulated FBP during starvation but failed to activate TORC1. We hypothesize that the production of FBP from glucose but not FBP itself is required for TORC1 activation. As the old adage goes ‘the journey is more important than the destination’ might be appropriate description for the canonical pathway of TORC1 activation. Determining how FBP activates TORC1 is an important challenge for the future.

### TORC1 is activated *via* the Rag GTPase-independent pathway

Our results support the notion that complete glycolysis is required for glucose-induced activation of TORC1 *via* the Gtr1-independent pathway. Blocking glycolysis *via* deleting/transcriptionally silencing genes that encode aldolase, triose-phosphate isomerase, phosphoglyceraldehyde dehydrogenase and phosphoglycerate kinase did not affect the Rag GTPase-dependent canonical pathway but abolished the Rag GTPase-independent pathway. Phenotypes of the *rav2Δ* cells and bafilomycin-treated *gtr1Δ* cells suggest that V-ATPase reassembly and V-ATPase activity are required for TORC1 activation *via* Rag GTPase-independent pathway.

Disassembly of V-ATPase during glucose starvation in yeast is a well-known phenomenon (Hayek et al., 2019). The V-ATPase consists of the membrane-bound Vo sector that forms the proton pore and the cytosolic V1 sector that has ATPase activity. V-ATPase catalyses ATP-dependent uptake of proton into the vacuole to regulates cytosolic pH. Glucose starvation results in dissociation of V1 from Vo and consequent inhibition of V-ATPase activity. Re-addition of glucose drives the reassembly of V-ATPase which is facilitated by the RAVE complex. We propose that vacuolar ATPase reassembly and activation following glucose metabolism acidifies the vacuole, which activates TORC1 resident in the vacuole. TORC1 activation in *gtr1Δ* cells is slower by about 10 minutes relative to wild type cells. Complete glycolysis and vacuolar acidification might take additional time which could account for the delay. Production of ATP *via* glycolysis could promote vacuolar ATPase reassembly and activity. Alternatively, metabolites derived from complete glycolysis could facilitate vacuolar ATPase reassembly. A previous study proposed that glucose regulates TORC1 by regulating the cytosolic pH via Gtr1 (Dechant et al., 2014). However, our results indicate that vacuolar ATPase regulates TORC1 independently of Gtr1. Addition of bafilomycin had very little effect on TORC1 activation on the canonical pathway and non-canonical pathways dependent on Rag GTPases.

### Glucose-induced TORC1 activation and AMPK inactivation are mechanistically independent

Addition of glucose activates TORC1 activation and AMPK inactivation. Our data indicate that these two events are not mechanistically dependent. Firstly, addition of 0.05% glucose/ 2-DG activates TORC1 but does not inactivate AMPK. Secondly, persistent activation of AMPK in *reg1* mutant cells does not affect TORC1 activation. Finally, rapamycin-mediated inhibition of TORC1 does not affect AMPK inactivation. However, our results are not consistent with results observed in other systems. AMPK negatively regulates TORC1 activity through multiple mechanisms in mammalian cells. In human cells, AMPK phosphorylates and activates TSC2, a negative regulator of mTORC1 (Inoki et al., 2003). AMPK also inhibits mTORC1 by phosphorylating its Raptor subunit (Gwinn et al., 2008). A more recent study showed a reciprocal antagonistic regulation of TORC1 and AMPK activities in fission yeast and mammalian cells (Ling et al., 2020). It is possible that AMPK and TORC1 are regulated differently in budding yeast compared to other organisms. But AMPK was also shown to inhibit TORC1 activity in budding yeast by phosphorylating the raptor subunit Kog1 (Hughes Hallett et al., 2015). However, we did not observe any difference between wild type Kog1 and its AMPK phosphosite mutant (Kog1^S491A,S494A^) in terms of glucose-induced TORC1 activation. This is unlikely to be due to the strain background as AMPK was shown not to play any in TORC1 inactivation during glucose starvation in another strain background (Prouteau et al., 2017).

In conclusion we identified three different mechanisms *via* glucose activates TORC1 (Figure 13). We show that G6P, FBP and complete glycolysis are required for non-canonical (Rag GTPase-dependent), canonical (Rag GTPase-dependent) and Rag GTPase-independent pathways, respectively. Glucose uptake and metabolism are often elevated in cancer cells (Hay, 2016). Cancer cells aerobically metabolise glucose and produce lactate even in the presence of mitochondria. This phenomenon of aerobic glycolysis is known as the ‘Warburg Effect’. However, how Warburg Effect benefits the growth and survival of cancer cells is not resolved. As glycolytic metabolites activate TORC1 by multiple mechanisms in yeast, it is possible that cancer cells might also use glycolysis to boost mTORC1 activity and positively regulate growth and proliferation. The glycolytic metabolite Dihydroxyacetone phosphate was shown to be sufficient to activate mTORC1 in human cells (Orozco et al., 2020). Interestingly, mTORC1 activity is found to be upregulated in several forms of cancer (Saxton and Sabatini, 2017). Uncovering the link between glycolysis and mTORC1 activity could identify novel targets for cancer therapy.

## Acknowledgments

We thank Prof. Claudio de Virgilio (University of Fribourg, Switzerland), Prof. Mike Hall (Biozentrum University of Basel, Switzerland), Prof. Capaldi (University of Arizona, USA) and Prof. Sheng-Cai Lin, Xiamen University, China). We would like to acknowledge A*STAR (Singapore)’s core funding provided to Arumugam laboratory.

## Author contributions

MA designed and performed TORC1 activation assay experiments, metabolite preparations, vacuolar ATPase reassembly experiments and contributed to writing the manuscript. LC performed the mass spectrometry experiments. JHCG, JHW and WJP performed TORC1 activation assays. JL performed the vacuolar ATPase reassembly experiments. PA#conceived and supervised the project and wrote the manuscript. All authors have read the manuscript.

### Declaration of interests

The authors declare no competing interests.

Please see the text for details

## METHODS

### Strains

All yeast strains used in this study were generated from *S. cerevisiae* SK1 genetic background. A complete list of strains and genotype information are provided in Table S1.

### TORC1 activity assay

Overnight cells were grown in medium at 30°C with 200 rpm shaking until they reached to logarithmic phase. Cells were then subjected to complete nutrient starvation by washing 3-times and incubating them in water at 30°C for 1 hour. Starved cells were then transferred to appropriate experimental solutions. Aliquots of the cultures were collected at different time points and used for preparing protein extracts. Phosphorylation of Sch9 was monitored by western blotting as described previously (Alfatah et al., 2021).

### Analysis of glycolytic metabolites

Glycolytic metabolites were extracted from logarithmic phase cells, starved cells and aliquots of cultures collected at different time points after the addition of carbon-source to starved cells. Cultures were quenched by mixing with twice the volume of methanol (pre-incubated at -80°C) and immediately spun down at 4500 rpm in centrifuge cooled to -10°C for 2 min. Supernatants were discarded and pellets were either stored at -20°C or processed further for metabolite extraction. Extraction solution, 40% (v/v) acetonitrile, 40% (v/v) methanol, and 20% (v/v) water was prepared and cooled to -80°C. All solvents used in extraction solution were HPLC grade. Extraction solution was mixed thoroughly prior to use. Pellets were resuspended with 700µl cold extraction solution by pipetting up and down and kept on ice for 15 min. Mixture was spun down at highest speed in centrifuge cooled to -4°C for 5 min to remove cell debris. Supernatants were transferred to 1.5 ml microcentrifuge tube and evaporated to dryness in a vacuum evaporator or stored at -80°C. The dried extracts were redissolved in 100 µl of 98:2 water/methanol and analyzed by targeted liquid chromatography-mass spectrometry (LC-MS) analysis as previously described (Zhong et al., 2017).

### Microscopy

Cultures at different time points were directly visualized under either bright-field or fluorescence microscope.

## REFERENCES

Alfatah, M., Wong, J.H., Krishnan, V.G., Lee, Y.C., Sin, Q.F., Goh, C.J.H., Kong, K.W., Lee, W.T., Lewis, J., Hoon, S., et al. (2021). TORC1 regulates the transcriptional response to glucose and developmental cycle via the Tap42-Sit4-Rrd1/2 pathway in Saccharomyces cerevisiae. BMC Biol 19, 95.

Binda, M., Bonfils, G., Panchaud, N., Peli-Gulli, M.P., and De Virgilio, C. (2010). An EGOcentric view of TORC1 signaling. Cell cycle 9, 221–222.

Bonfils, G., Jaquenoud, M., Bontron, S., Ostrowicz, C., Ungermann, C., and De Virgilio, C. (2012). Leucyl-tRNA synthetase controls TORC1 via the EGO complex. Molecular cell 46, 105–110.

Bowman, E.J., Siebers, A., and Altendorf, K. (1988). Bafilomycins: a class of inhibitors of membrane ATPases from microorganisms, animal cells, and plant cells. Proceedings of the National Academy of Sciences of the United States of America 85, 7972–7976.

Chantranupong, L., Wolfson, R.L., Orozco, J.M., Saxton, R.A., Scaria, S.M., Bar-Peled, L., Spooner, E., Isasa, M., Gygi, S.P., and Sabatini, D.M. (2014). The Sestrins interact with GATOR2 to negatively regulate the amino-acid-sensing pathway upstream of mTORC1. Cell reports 9, 1–8.

Conrad, M., Schothorst, J., Kankipati, H.N., Van Zeebroeck, G., Rubio-Texeira, M., and Thevelein, J.M. (2014). Nutrient sensing and signaling in the yeast Saccharomyces cerevisiae. FEMS Microbiol Rev 38, 254–299.

Dechant, R., Saad, S., Ibanez, A.J., and Peter, M. (2014). Cytosolic pH regulates cell growth through distinct GTPases, Arf1 and Gtr1, to promote Ras/PKA and TORC1 activity. Molecular cell 55, 409–421.

Fenton, A.W., and Blair, J.B. (2002). Kinetic and allosteric consequences of mutations in the subunit and domain interfaces and the allosteric site of yeast pyruvate kinase. Arch Biochem Biophys 397, 28–39.

Gonzalez, A., and Hall, M.N. (2017). Nutrient sensing and TOR signaling in yeast and mammals. The EMBO journal 36, 397–408.

Gwinn, D.M., Shackelford, D.B., Egan, D.F., Mihaylova, M.M., Mery, A., Vasquez, D.S., Turk, B.E., and Shaw, R.J. (2008). AMPK phosphorylation of raptor mediates a metabolic checkpoint. Molecular cell 30, 214–226.

Hay, N. (2016). Reprogramming glucose metabolism in cancer: can it be exploited for cancer therapy? Nat Rev Cancer 16, 635–649.

Hayek, S.R., Rane, H.S., and Parra, K.J. (2019). Reciprocal Regulation of V-ATPase and Glycolytic Pathway Elements in Health and Disease. Front Physiol 10, 127.

Herzig, S., and Shaw, R.J. (2018). AMPK: guardian of metabolism and mitochondrial homeostasis. Nature reviews. Molecular cell biology 19, 121–135.

Hughes Hallett, J.E., Luo, X., and Capaldi, A.P. (2014). State transitions in the TORC1 signaling pathway and information processing in Saccharomyces cerevisiae. Genetics 198, 773–786.

Hughes Hallett, J.E., Luo, X., and Capaldi, A.P. (2015). Snf1/AMPK promotes the formation of Kog1/Raptor-bodies to increase the activation threshold of TORC1 in budding yeast. eLife 4.

Inoki, K., Zhu, T., and Guan, K.L. (2003). TSC2 mediates cellular energy response to control cell growth and survival. Cell 115, 577–590.

Jurica, M.S., Mesecar, A., Heath, P.J., Shi, W., Nowak, T., and Stoddard, B.L. (1998). The allosteric regulation of pyruvate kinase by fructose-1,6-bisphosphate. Structure 6, 195–210.

Kane, P.M., and Smardon, A.M. (2003). Assembly and regulation of the yeast vacuolar H+-ATPase. J Bioenerg Biomembr 35, 313–321.

Kondo, T., and Beutler, E. (1979). Depletion of red cell ATP by incubation with 2-deoxyglucose. J Lab Clin Med 94, 617–623.

Ling, N.X.Y., Kaczmarek, A., Hoque, A., Davie, E., Ngoei, K.R.W., Morrison, K.R., Smiles, W.J., Forte, G.M., Wang, T., Lie, S., et al. (2020). mTORC1 directly inhibits AMPK to promote cell proliferation under nutrient stress. Nat Metab 2, 41–49.

Marsh, J.J., and Lebherz, H.G. (1992). Fructose-bisphosphate aldolases: an evolutionary history. Trends in biochemical sciences 17, 110–113.

Nicastro, R., Sardu, A., Panchaud, N., and De Virgilio, C. (2017). The Architecture of the Rag GTPase Signaling Network. Biomolecules 7.

Orozco, J.M., Krawczyk, P.A., Scaria, S.M., Cangelosi, A.L., Chan, S.H., Kunchok, T., Lewis, C.A., and Sabatini, D.M. (2020). Dihydroxyacetone phosphate signals glucose availability to mTORC1. Nat Metab 2, 893–901.

Panchaud, N., Peli-Gulli, M.P., and De Virgilio, C. (2013). SEACing the GAP that nEGOCiates TORC1 activation: evolutionary conservation of Rag GTPase regulation. Cell cycle 12, 2948–2952.

Peli-Gulli, M.P., Sardu, A., Panchaud, N., Raucci, S., and De Virgilio, C. (2015). Amino Acids Stimulate TORC1 through Lst4-Lst7, a GTPase-Activating Protein Complex for the Rag Family GTPase Gtr2. Cell reports 13, 1–7.

Powis, K., and De Virgilio, C. (2016). Conserved regulators of Rag GTPases orchestrate amino acid-dependent TORC1 signaling. Cell discovery 2, 15049.

Prouteau, M., Desfosses, A., Sieben, C., Bourgoint, C., Lydia Mozaffari, N., Demurtas, D., Mitra, A.K., Guichard, P., Manley, S., and Loewith, R. (2017). TORC1 organized in inhibited domains (TOROIDs) regulate TORC1 activity. Nature 550, 265–269.

Roberts, D.J., Tan-Sah, V.P., Ding, E.Y., Smith, J.M., and Miyamoto, S. (2014). Hexokinase-II positively regulates glucose starvation-induced autophagy through TORC1 inhibition. Molecular cell 53, 521–533.

Saxton, R.A., Knockenhauer, K.E., Schwartz, T.U., and Sabatini, D.M. (2016a). The apo-structure of the leucine sensor Sestrin2 is still elusive. Science signaling 9, ra92.

Saxton, R.A., Knockenhauer, K.E., Wolfson, R.L., Chantranupong, L., Pacold, M.E., Wang, T., Schwartz, T.U., and Sabatini, D.M. (2016b). Structural basis for leucine sensing by the Sestrin2-mTORC1 pathway. Science 351, 53–58.

Saxton, R.A., and Sabatini, D.M. (2017). mTOR Signaling in Growth, Metabolism, and Disease. Cell 168, 960–976.

Slater, E.C. (1973). The mechanism of action of the respiratory inhibitor, antimycin. Biochim Biophys Acta 301, 129–154.

Smardon, A.M., Diab, H.I., Tarsio, M., Diakov, T.T., Nasab, N.D., West, R.W., and Kane, P.M. (2014). The RAVE complex is an isoform-specific V-ATPase assembly factor in yeast. Molecular biology of the cell 25, 356–367.

Smardon, A.M., Tarsio, M., and Kane, P.M. (2002). The RAVE complex is essential for stable assembly of the yeast V-ATPase. The Journal of biological chemistry 277, 13831–13839.

Urban, J., Soulard, A., Huber, A., Lippman, S., Mukhopadhyay, D., Deloche, O., Wanke, V., Anrather, D., Ammerer, G., Riezman, H., et al. (2007). Sch9 is a major target of TORC1 in Saccharomyces cerevisiae. Molecular cell 26, 663–674.

Wolfson, R.L., Chantranupong, L., Saxton, R.A., Shen, K., Scaria, S.M., Cantor, J.R., and Sabatini, D.M. (2016). Sestrin2 is a leucine sensor for the mTORC1 pathway. Science 351, 43–48.

Xu, Y.F., Zhao, X., Glass, D.S., Absalan, F., Perlman, D.H., Broach, J.R., and Rabinowitz, J.D. (2012). Regulation of yeast pyruvate kinase by ultrasensitive allostery independent of phosphorylation. Molecular cell 48, 52–62.

Zhang, C.S., Hawley, S.A., Zong, Y., Li, M., Wang, Z., Gray, A., Ma, T., Cui, J., Feng, J.W., Zhu, M., et al. (2017). Fructose-1,6-bisphosphate and aldolase mediate glucose sensing by AMPK. Nature 548, 112–116.

Zhong, W., Cui, L., Goh, B.C., Cai, Q., Ho, P., Chionh, Y.H., Yuan, M., Sahili, A.E., Fothergill-Gilmore, L.A., Walkinshaw, M.D., et al. (2017). Allosteric pyruvate kinase-based “logic gate” synergistically senses energy and sugar levels in Mycobacterium tuberculosis. Nature communications 8, 1986.

