## Supplementary data for "Metabolism of glucose activates TORC1 through multiple mechanisms in *Saccharomyces cerevisiae*"

**Supplementary figure legends**

**Figure S1 2-deoxyglucose (2-DG) activates TORC1**

A. Wild type cells in logarithmic phase (C) in SC-Glu medium were subjected to complete nutrient starvation by incubating them in water for 1 hour. Starved cells (S) were then transferred to a solution containing 2% glucose or 2% 2-deoxyglucose or 2% 6-deoxyglucose in the presence and absence of rapamycin (2 µM). Aliquots of the cultures were taken after 0', 10', 20' and 30' and were used for preparing protein extracts. Phosphorylation of Sch9 was monitored by Western blotting.
B. Same as A but the assay was performed with different concentrations of 2-deoxyglucose.

**Figure S2 Glycolysis but not the Pentose Phosphate pathway (PPP) is required for glucose-induced TORC1 activation via the Rag GTPase-independent pathway**

A. Wild type, *gtr1Δ, zwf1Δ* and *zwf1Δ gtr1Δ* cells were grown to logarithmic phase (C) in SC-Glu medium and were subjected to complete nutrient starvation by incubating them in water for 1 hour. Starved cells (S) were then transferred to a solution containing 2% glucose. Aliquots of the cultures taken after 0', 10', 20' and 30' were used for preparing protein extracts. Phosphorylation of Sch9 was monitored by Western blotting.
B. Same as A but performed with wild type, *gtr1Δ, pfk1Δ* and *pfk1Δ gtr1Δ* cells.

**Figure S3 2-DG mediated TORC1 activation is positively and negatively regulated by SEACIT and Lst4**

A. Wild type, *npr2Δ, iml1Δ* and *npr2Δ gtr1Δ* and *iml1Δ gtr1Δ* cells were grown to logarithmic phase (C) in SC-Glu medium and were subjected to complete nutrient starvation by incubating them in water for 1 hour. Starved cells (S) were then transferred to either a solution containing 2% glucose or 0.05% 2-DG. Aliquots of the cultures taken after 0', 10', 20' and 30' were used for preparing protein extracts. Phosphorylation of Sch9 was monitored by Western blotting.

B. Wild type, *gtr1Δ, npr3Δ* and *lst4Δ* cells were grown to logarithmic phase (C) in SC-Glu medium and were subjected to complete nutrient starvation by incubating them in water for 1 hour. Starved cells (S) were then transferred to either a solution containing 2% glucose or 0.05% 2-DG. Aliquots of the cultures taken after 0', 10', 20' and 30' were used for preparing protein extracts. Phosphorylation of Sch9 was monitored by Western blotting.

**Figure S4 Like glucose 6-phosphate, mannose 6-phosphate can activate the non-canonical Rag GTPase-dependent pathway of TORC1 activation**

Wild type, *gtr1Δ, pmi40Δ* and *pmi40Δ gtr1Δ* cells were grown to logarithmic phase (C) in SC-Glu supplemented with 5 mM mannose and were subjected to complete nutrient starvation by incubating them in water for 1 hour. Starved cells (S) were then transferred to a solution containing 2% glucose or 2% mannose or 0.05% mannose. Aliquots of the cultures taken after 0', 10', 20' and 30' were used for preparing protein extracts. Phosphorylation of Sch9 was monitored by Western blotting.

**Figure S5 Analyses of glycolytic metabolites in wild type and *tpi1Δ* cells subjected to the TORC1 activation assay**

Wild type and *tpi1Δ* cells were grown in SC-EtOH/Gly medium and then subjected to complete nutrient starvation by incubating them in water for 1 hour. Starved cells (S) were transferred to a solution containing 2% glucose. Aliquots of the cultures were taken after 0', 10', 20' and 30' for extracting metabolites for LC-MS analysis. Relative molar levels of various glycolytic metabolites in wild type and *tpi1Δ* cells at different stages of glucose-induced TORC1 activation assay are shown.

**Figure S6 Analyses of glycolytic metabolites in wild type, *pgk1Δ*, *cdc19*Δ and *pgm1*Δ cells subjected to the TORC1 activation assay**

A. Wild type, *pgk1Δ*, *cdc19Δ* and *pgm1Δ* cells were grown in SC-EtOH/Gly medium and then subjected to complete nutrient starvation by incubating them in water for 1 hour. Starved cells (S) were transferred to a solution containing 2% glucose. Aliquots of the cultures were taken after 0', 10', 20' and 30' for extracting metabolites for LC-MS analysis. Relative molar levels of various glycolytic metabolites in wild type *cdc19Δ* and *pgm1Δ* cells at different stages of glucose-induced TORC1 activation assay are shown.

B. Wild type, *cdc19Δ, pgm1Δ*, *gtr1Δ*, *cdc19Δ gtr1Δ* and *pgm1Δ gtr1Δ* cells were grown in SC-EtOH/Gly medium and then subjected to complete nutrient starvation by incubating them in water for 1 hour. Starved cells (S) were transferred to a solution containing 2% glucose. Sch9 phosphorylation was assayed by Western blotting.

**Figure S7 FBP-binding proteins and the canonical pathway**

A. Wild type, *npr3Δ*, *gtr1Δ* and *npr3Δ gtr1Δ* cells expressing native aldolase or human aldolase A were grown in logarithmic phase (C) in SC-Glu medium were subjected to complete nutrient starvation by incubating them in water for 1 hour. Starved cells (S) were then transferred to a solution containing 2% glucose. Aliquots of the cultures were taken after 0’, 10’, 20’ and 30’ and were used for preparing protein extracts. Phosphorylation of Sch9 was monitored by Western blotting.

B. Similar to A but performed with wild type, *gtr1Δ*, *npr3Δ*, *fbp1Δ*, *fbp1Δ gtr1Δ* and *fbp1Δ npr3Δ* cells

C. Similar to A but performed with Wild type, *npr3Δ*, *gtr1Δ* cells expressing either $Cdc19^{WT}$ or $Cdc19^{E392A\ R459Q}$.

#

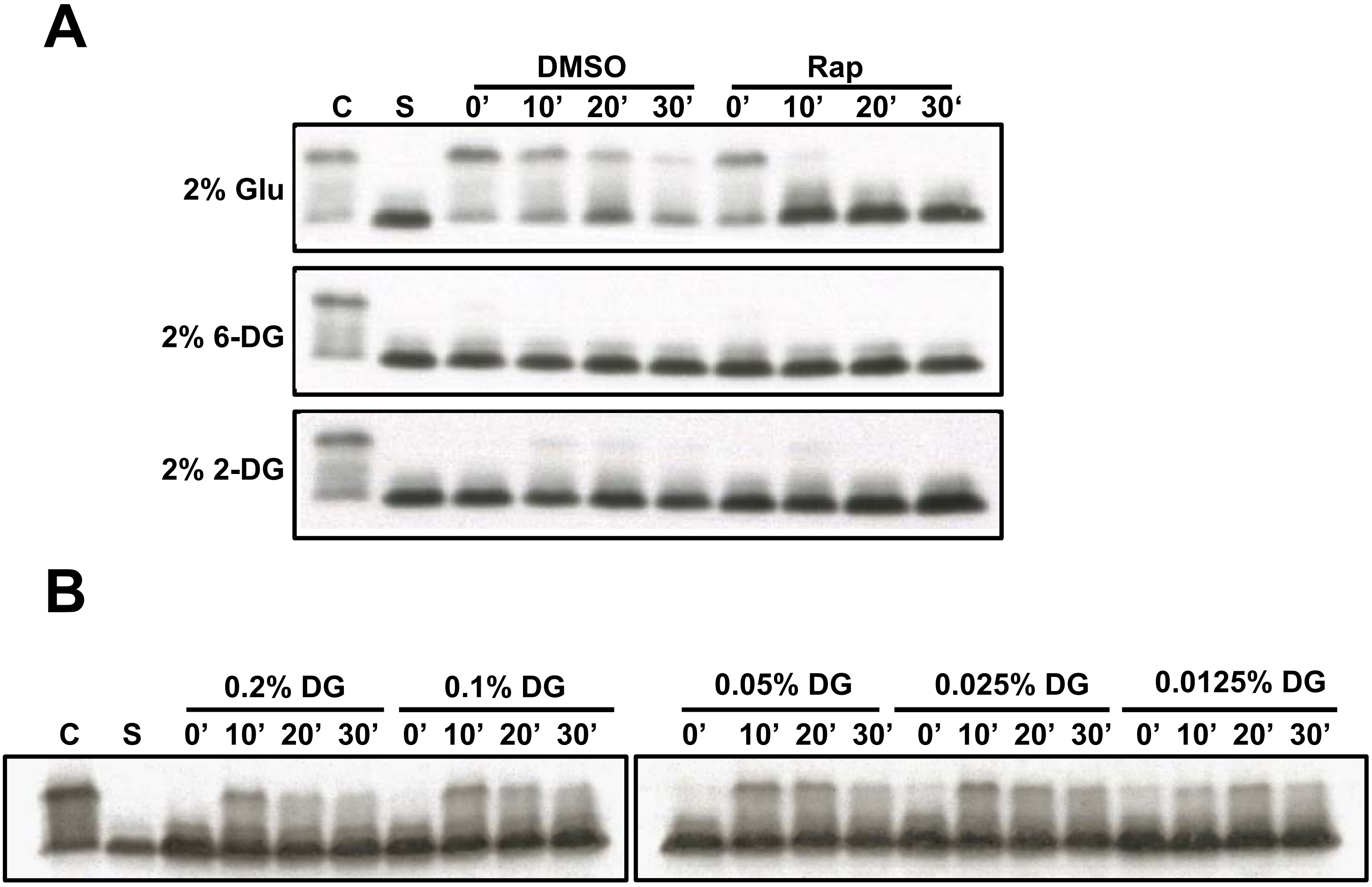
A
DMSO
Rap
C S 0' 10' 20' 30' 0' 10' 20' 30'
2% Glu
2% 6-DG
2% 2-DG
B
0.2% DG
0.1% DG
C S 0' 10' 20' 30' 0' 10' 20' 30'
0.05% DG
0.025% DG
0.0125% DG
0' 10' 20' 30' 0' 10' 20' 30' 0' 10' 20' 30'

### Figure S1

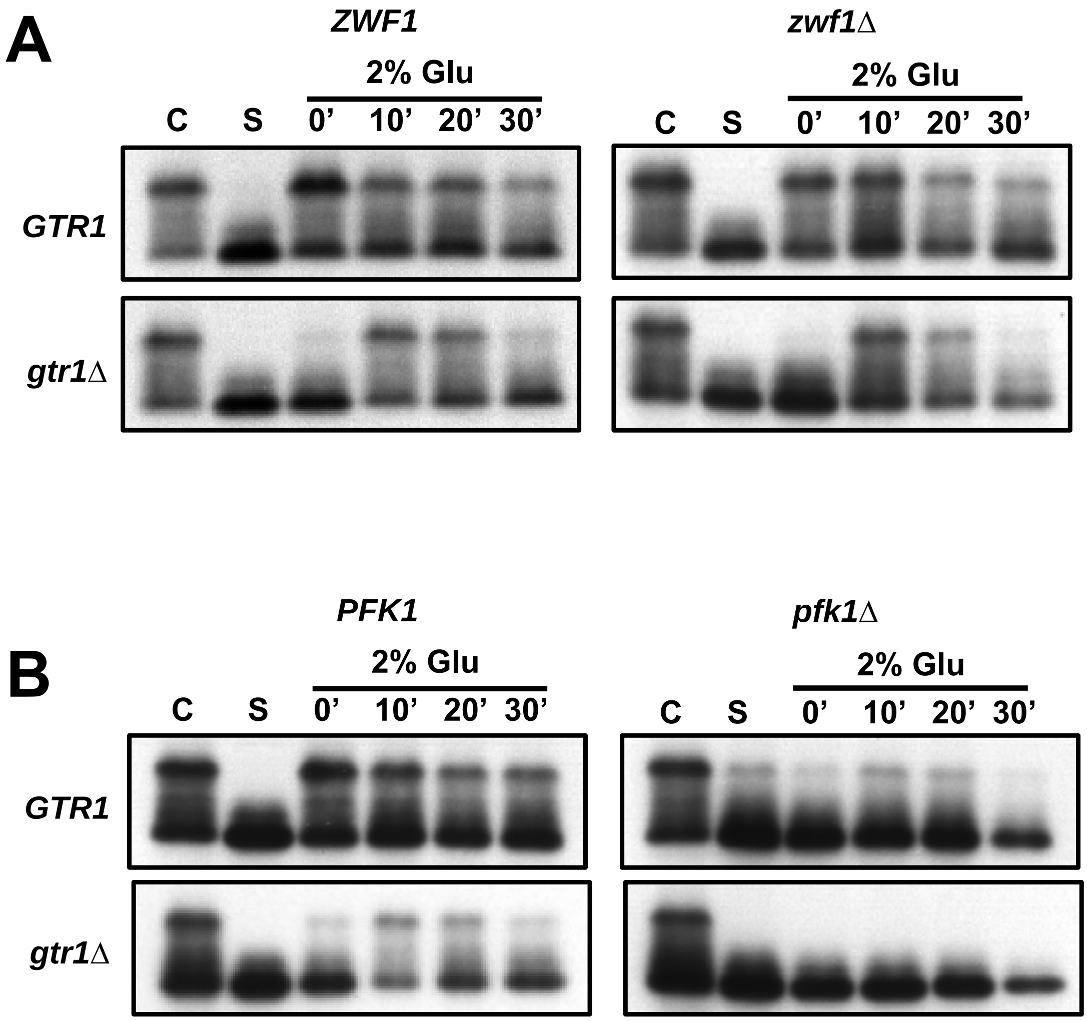


### Figure S2

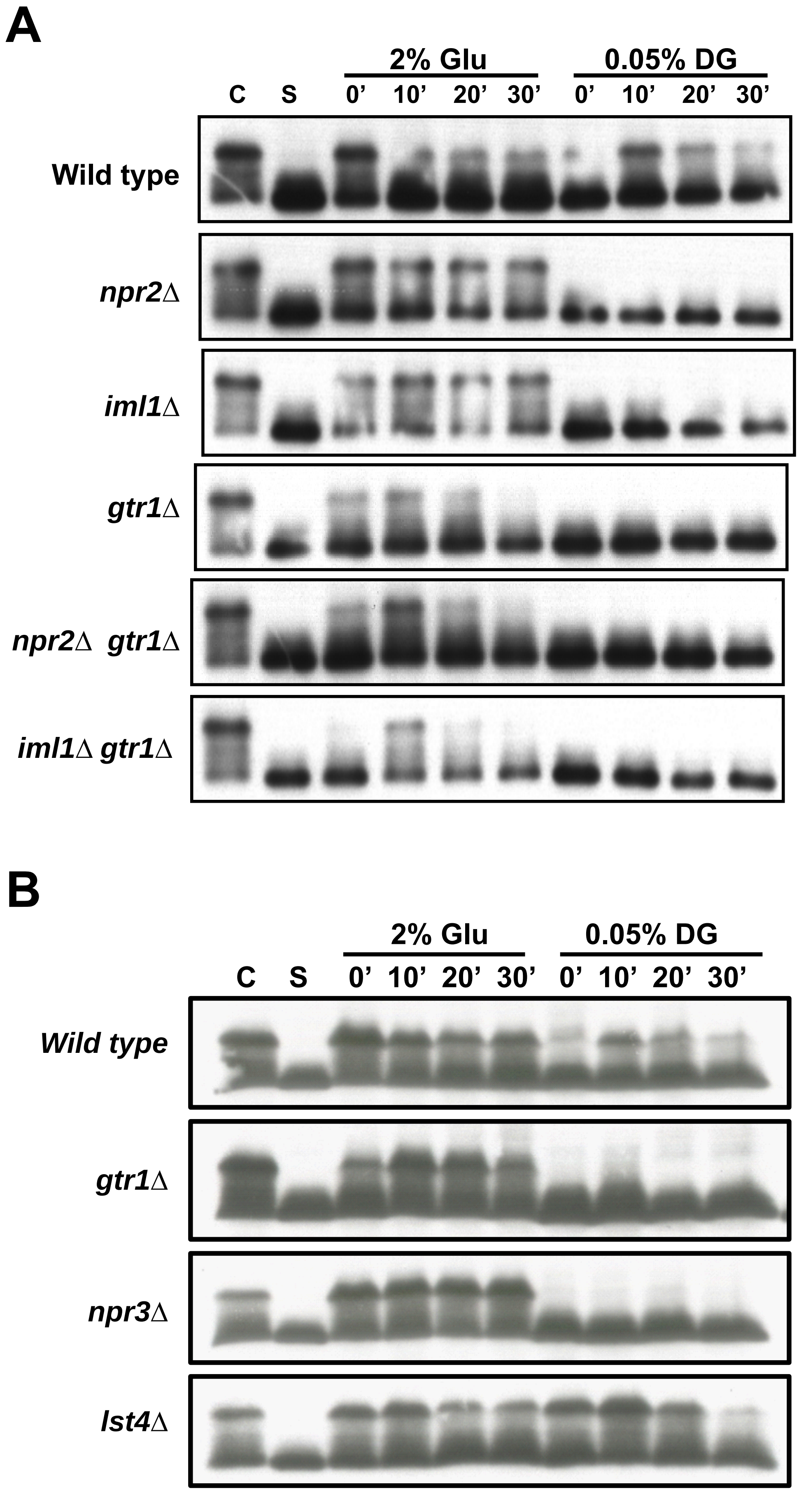
A
2% Glu
0.05% DG
C
S
0'
10'
20'
30'
0'
10'
20'
30'
Wild type
npr2Δ
iml1Δ
gtr1Δ
npr2Δ gtr1Δ
iml1Δ gtr1Δ
B
2% Glu
0.05% DG
C
S
0'
10'
20'
30'
0'
10'
20'
30'
Wild type
gtr1Δ
npr3Δ
lst4Δ

### Figure S3

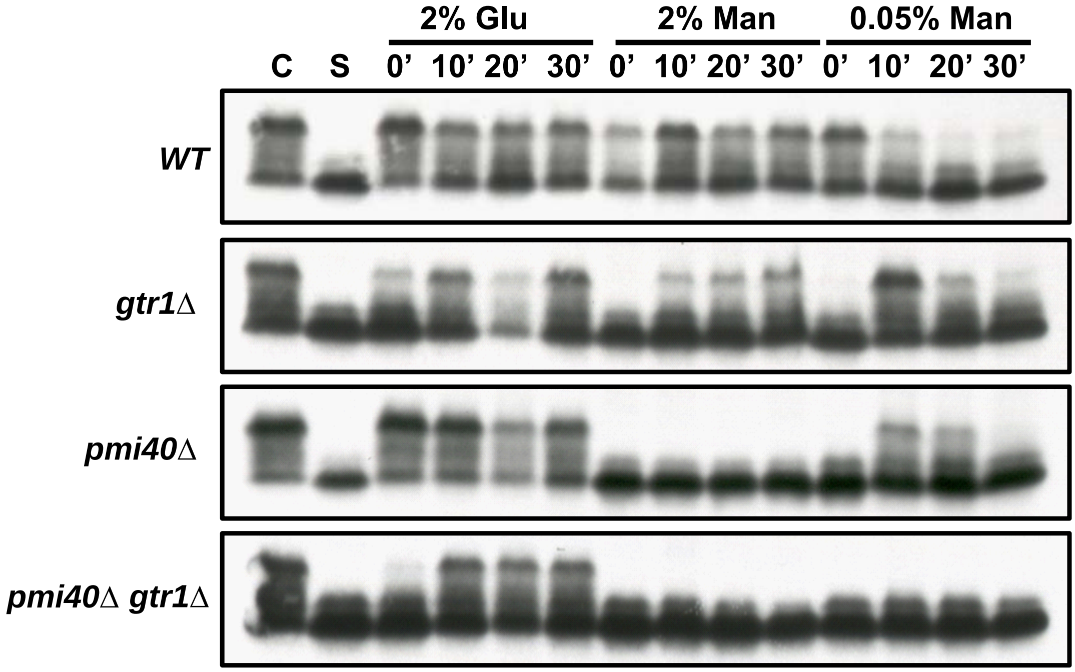
2% Glu
2% Man
0.05% Man
C S 0' 10' 20' 30' 0' 10' 20' 30' 0' 10' 20' 30'
WT
gtr1Δ
pmi40Δ
pmi40Δ gtr1Δ

### Figure S4

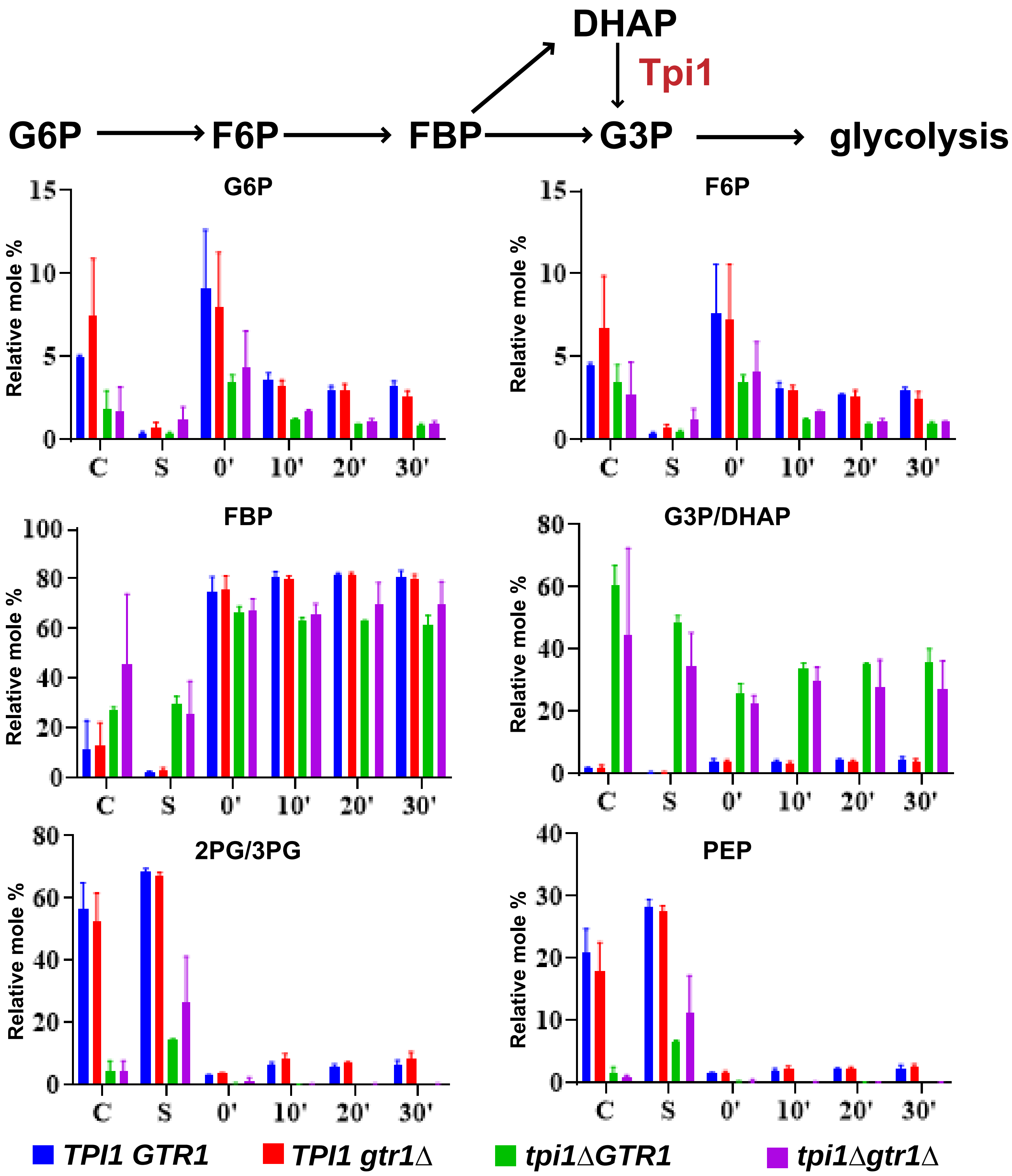
DHAP
Tpi1
G6P
F6P
FBP
G3P
glycolysis
G6P
F6P
FBP
G3P/DHAP
2PG/3PG
PEP
Relative mole %
C
S
0'
10'
20'
30'
TPI1 GTR1
TPI1 gtr1Δ
tpi1ΔGTR1
tpi1Δgtr1Δ

### Figure S5

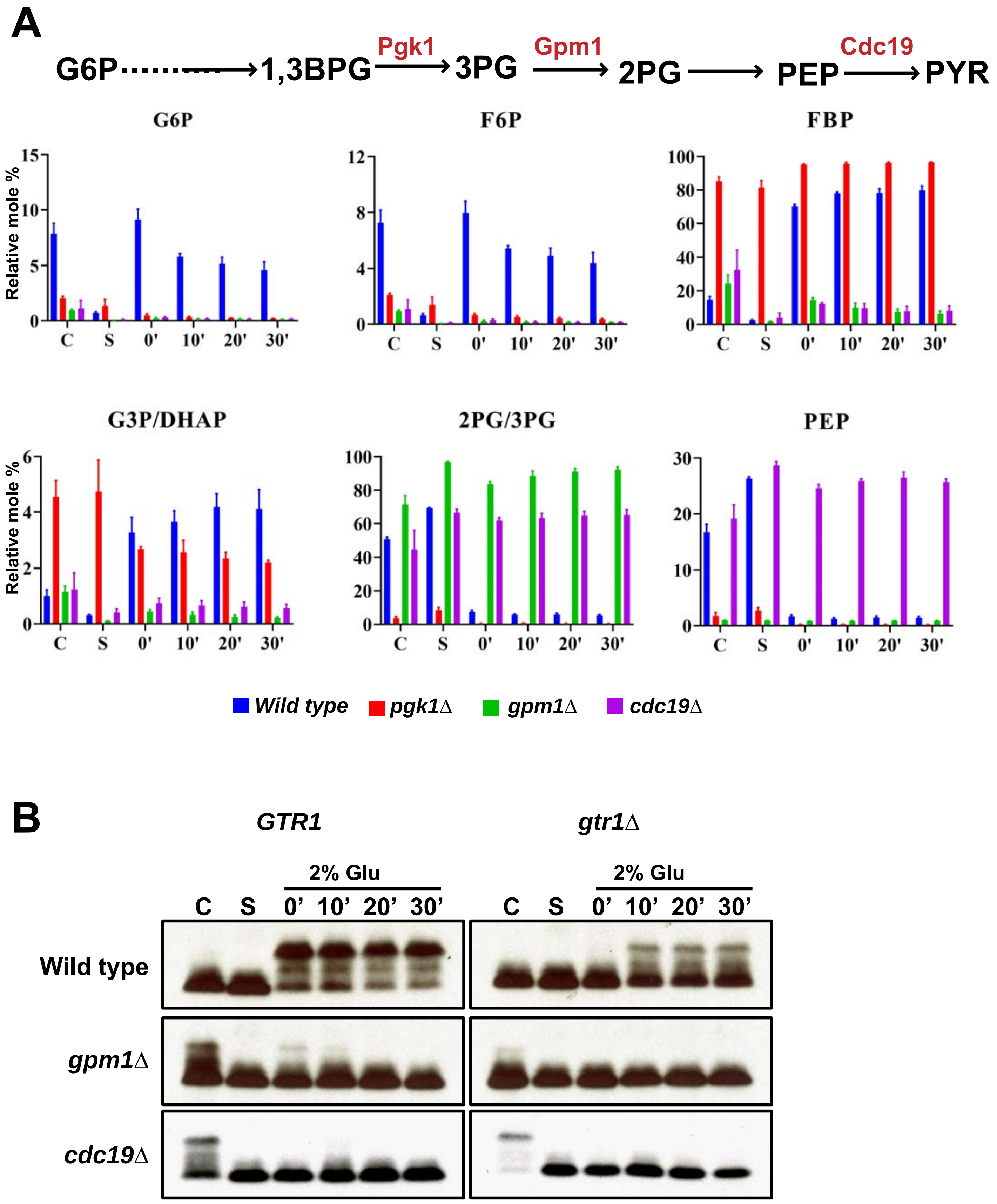
A
G6P
1,3BPG
Pgk1
3PG
Gpm1
2PG
PEP
Cdc19
PYR
G6P
F6P
FBP
G3P/DHAP
2PG/3PG
PEP
Relative mole %
C
S
0'
10'
20'
30'
Wild type
pgk1Δ
gpm1Δ
cdc19Δ
B
GTR1
gtr1Δ
2% Glu
Wild type
gpm1Δ
cdc19Δ

### Figure S6

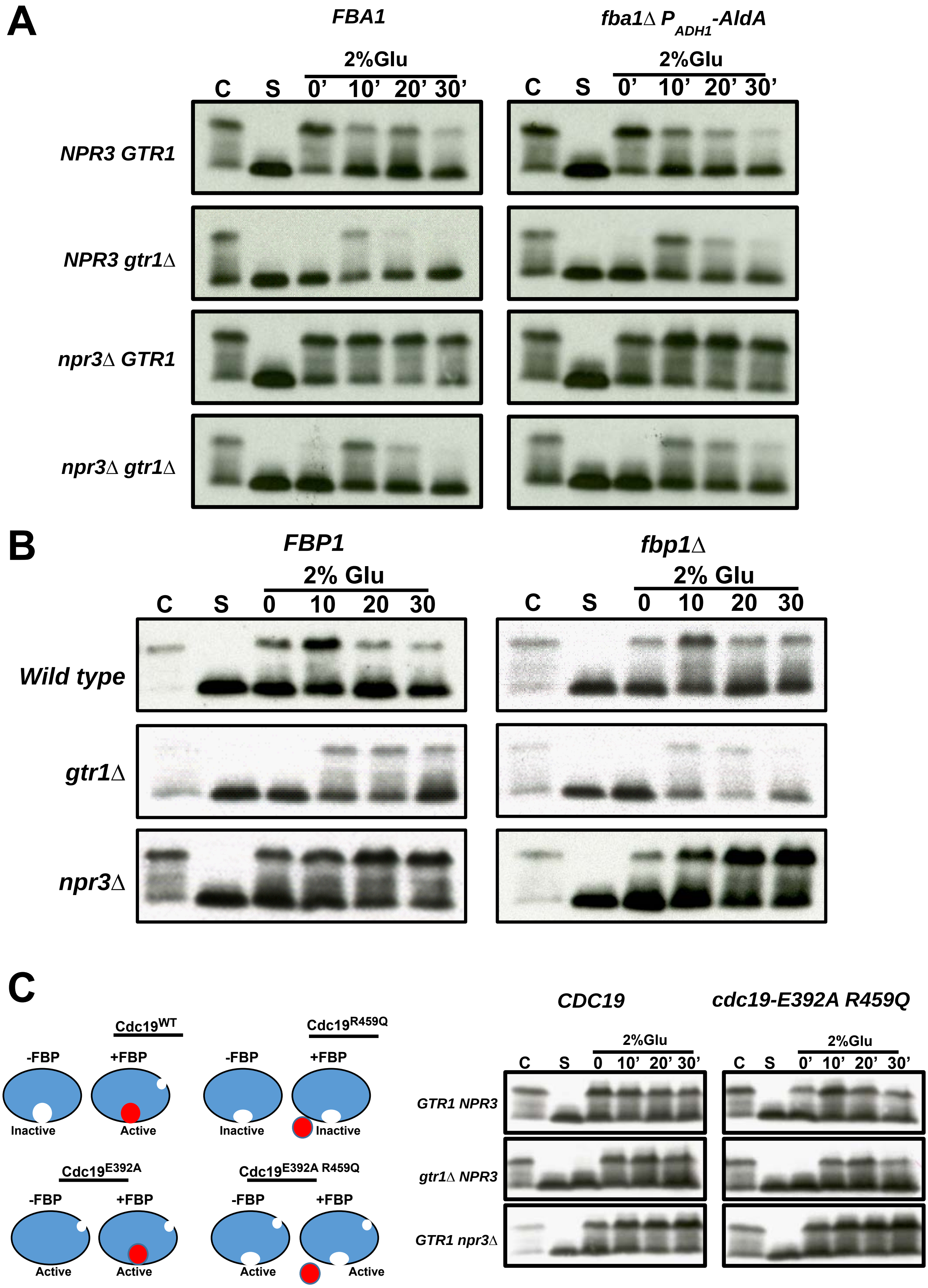
A
FBA1
fba1Δ P ADH1-AldA
2%Glu
C S 0’ 10’ 20’ 30’
2%Glu
C S 0’ 10’ 20’ 30’
NPR3 GTR1
NPR3 gtr1Δ
npr3Δ GTR1
npr3Δ gtr1Δ
B
FBP1
fbp1Δ
2% Glu
C S 0 10 20 30
2% Glu
C S 0 10 20 30
Wild type
gtr1Δ
npr3Δ
C
Cdc19WT
-FBP
+FBP
Inactive
Active
Cdc19R459Q
-FBP
+FBP
Inactive
Inactive
Cdc19E392A
-FBP
+FBP
Active
Active
Cdc19E392A R459Q
-FBP
+FBP
Active
Active
CDC19
cdc19-E392A R459Q
2%Glu
C S 0 10’ 20’ 30’
2%Glu
C S 0’ 10’ 20’ 30’
GTR1 NPR3
gtr1Δ NPR3
GTR1 npr3Δ

### Figure S7
